# DETECTION OF SURFACE FORCES BY A CELL WALL MECHANOSENSOR

**DOI:** 10.1101/2020.12.10.418905

**Authors:** Ramakanth Neeli-Venkata, Ruben Celador, Yolanda Sanchez, Nicolas Minc

## Abstract

Surface receptors of animal cells, such as integrins, promote mechanosensation by forming local clusters as signaling hubs that transduce tensile forces. Walled cells of plants and fungi also feature surface sensors, with long extracellular domains embedded in their cell wall (CW), thought to detect CW injuries and promote repair. How these sensors probe surface forces remains unknown. By studying the conserved CW sensor Wsc1 in fission yeast, we uncovered the formation of micrometer-sized clusters at sites of local force application onto the CW. These clusters form within minutes of CW compression, in dose-dependence with mechanical stress and dissolve upon stress relaxation. Our data support that Wsc1 senses CW mechanical stress and accumulates to local sites of enhanced stress through its CW-associated extracellular WSC domain, independently of canonical polarity, trafficking and downstream CW regulatory pathways. Wsc1 may represent an autonomous module to detect and transduce local surface forces onto the CW.

## INTRODUCTION

Cells experience a large range of tensile, compressive or shear forces from their environment, that influence growth, migration or differentiation (Vogel and Sheetz, 2006). In animal cells, forces are typically detected at the cell surface through specific classes of integral membrane receptors, such as integrins or cadherins, which interact with the extracellular matrix or neighbouring cells (Kechagia, et al., 2019; Ladoux and Mège, 2017). Those receptors are connected to downstream elements; including scaffolding proteins, kinases and phosphatases, and cytoskeletal components, that together allow forces to be detected, transmitted and transduced into biochemical signals that initiate adaptive responses (Chen, et al., 2017). One conserved feature of surface mechanosensing stands in the formation of receptor clusters that maturate into larger assemblies in response to tensile forces (Galbraith, et al., 2002). Clusters may have sizes ranging from few tens of nanometers up to micrometers, and contain hundreds to thousands of molecules (Changede and Sheetz, 2017). Their formation and maturation under force could implicate cis-cis interactions, reduced local diffusivity, or membrane trafficking pathways, all dependent on local surface forces (Moreno-Layseca, et al., 2019; Welf, et al., 2012; Paszek, et al., 2009). The functional role of clustering includes larger force-bearing and detection, noise-reduction in mechano-perception, as well as the separation of sensing units that probe local forces around cells at an intermediate scale much larger than that of single molecules (Changede and Sheetz, 2017). Thus, surface sensor clustering may constitute a generic modular feature for cellular mechanobiology.

Non-motile walled cells, such as plant and fungal cells, also express surface sensors which may primarily interact and monitor properties of their cell walls (CW) (Pérez, et al., 2018; Hamann and Denness, 2011; Levin, 2011). Yeast and fungi, for instance, feature two conserved families of single-pass transmembrane CW sensors of the WSC-type and MID-type. In analogy to animal receptors, these sensors exhibit long ∼50-80nm extracellular domains embedded into the CW that have been proposed to probe surface stress and integrity (Elhasi and Blomberg, 2019; Kock, et al., 2015). They function as upstream trigger of the Cell Wall Integrity pathway (CWI), and activate CW associated Rho GTPases, and/or the expression of CW repair genes through protein kinases C and the Pmk1 MAPK, in response to global stresses such as those caused by e.g. heat or antifungal agents (Pérez, et al., 2018; Levin, 2011). Sensors from both families share a similar architecture with a cytoplasmic C-terminal tail that mediate downstream signaling, followed by a single trans<u>m</u>embrane <u>d</u>omain (TMD), an O-mannosylated <u>s</u>erine/threonine-rich region (STR), a head group and a signal sequence. The head group of the WSC-type sensors is a conserved WSC cysteine-rich domain, and that of the MID-type is an N-glycosylated asparagine. These head groups are thought to interact with CW polysaccharides. For instance, WSC domains belong to C-type lectin domain families that bind carbohydrates plausibly through multiple weak interactions (Kock, et al., 2015). Pioneering studies using Atomic Force Microscopy (AFM) on engineered polypeptides made of *S. cerevisiae* Wsc1 and Mid2 STR demonstrated that these domains can behave as nano-springs (expanding linearly with force) (Dupres, et al., 2009). Therefore, CW sensors may embrace functional domains to interact with the CW, sense surface forces and transduce them downstream. Accordingly, recent reports have directly implicated these sensors in CW mechanosensation and homeostasis (Banavar, et al., 2018; Davì, et al., 2018; Mishra, et al., 2017). To date, however, whether and how CW nano-sensors can directly probe CW mechanics or surface forces remain unknown (Kock, et al., 2015).

We here report on the formation of large and stable micrometer-sized Wsc1 clusters at local sites of cell-cell contacts and CW compression in the fission yeast *Schizosaccarhomyces pombe*. We establish the dose-dependence of cluster formation on CW mechanical stress, their dynamics of assembly and disassembly in response to compression and relaxation respectively, and their function in supporting survival in packed environments. Our findings suggest that Wsc1 probe surface mechanical stress and clusters at sites of enhanced stress through its extracellular WSC domain, independently of polarity and trafficking regulators and other canonical elements of the CWI pathway. We propose that Wsc1 may represent an autonomous sensing module for detecting and transducing local surface compressive forces on the CW.

## RESULTS

### Wsc1 forms clusters at sites of cell-cell contacts

Fission yeast features only two CW sensors, Wsc1 and Mtl2, which complement each other to support cell viability. Mtl2-GFP is localized all around the cell surface, while Wsc1-GFP is enriched at cell tips and also to the septum during cell division (Cruz, et al., 2013). As previously reported in *S. cerevisiae*, we observed that Wsc1-GFP distribution around cell tips was non-uniform, with the clear appearance of little foci/clusters enriched in Wsc1 (Figure S1A) (Kock, et al., 2015). Remarkably, by filming cells growing and dividing in standard agar pads, we made the serendipitous observation of the transient formation of bright and large micrometer-sized Wsc1-GFP clusters at sites of new end-new end contacts after cell division (Figure 1A, and Movies S1-S2). These clusters typically formed within few tens of minutes after the completion of septation, when new ends round up and push onto each other due to turgor forces (Atilgan, et al., 2015). Clusters were more prominent and stable in time when cells could not grow away from each other due to the presence of neighbors. In addition, upon cell rearrangement, cells frequently slid past each other, relaxing the contact and causing cluster disappearance within few minutes (Figure 1A-1C, and Movie S2). Local Wsc1-GFP clusters, with similar appearance and disappearance kinetics were also often observed when cells grew onto each other and contacted old end against old end or old end against cell sides, suggesting that they are not specific to septation (Figure 1D-1E, and Movies S3). Finally, in several instances of cells contacting and sliding, Wsc1 clusters appeared to track the site of cell contact (Figure S1B, Movies S2). These observations suggest that Wsc1-GFP may dynamically sense and cluster at firm cell-cell contacts.

**Figure 1.**
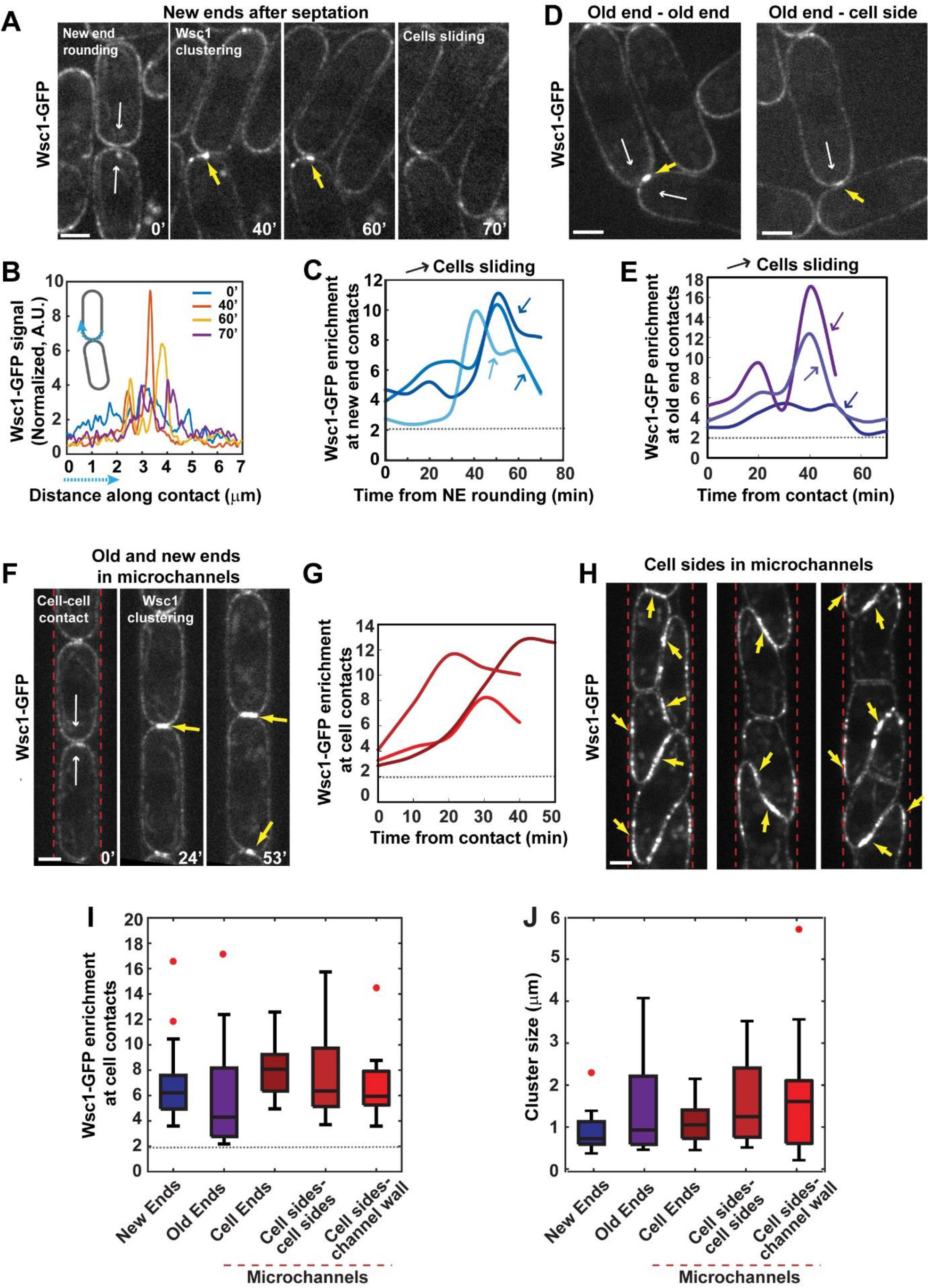
Wsc1 surface sensor forms micrometer-sized clusters at sites of cell-cell contacts. **(A)** Spinning disk confocal mid-plane time-lapse of wild-type cells expressing Wsc1-GFP growing and dividing on standard agar pads. Note the appearance of bright clusters at the contact between new ends after septation (40’ and 60’), which disappear when cells slide apart and release pressure on the contact (70’). White arrows depict the direction of pressure-derived forces on the contact, and yellow arrows point to Wsc1 clusters. (**B)** Normalized fluorescence levels of Wsc1-GFP along the new end contact (blue dotted line in the inset), at each time point from the example from 1A. **(C)** Dynamic of Wsc1-GFP enrichment in three representative pairs of dividing cells. Enrichment is calculated as the ratio between the maximum fluorescent signal at the site of contact to that on cell sides with no contact. The colored arrows indicate the moment when cells slide apart. Dotted line in the graph marks the reference value of enrichment of 2, corresponding to the presence of two apposed membranes. **(D)** Wsc1-GFP clusters formed at an old end-old end or old end-cell side contact. **(E)** Wsc1-GFP enrichment dynamics at old end contacts in three representative cell pairs. **(F)** Time-lapse spinning disk confocal mid-section of wild-type cells expressing Wsc1-GFP grown in linear microfabricated channels, demarked by red dotted lines. **(G)** Wsc1-GFP enrichment dynamics at old or new end contacts in three representative cell pairs grown in microchannels. **(H)** Examples of 3 individual microchannels in which cells have grown for longer periods of times (∼5h), adopting complex morphologies and packed arrangements, resulting in Wsc1-GFP decorating large parts of cell sides even those pressed against channel walls. **(I-J)** Box plots with outliers as red dots of Wsc1-GFP enrichment and cluster sizes in the different conditions presented in 1A-1H. Cluster size is calculated as the full width at mid-height of a Gaussian fit of Wsc1 signal along the contact. (n= 16, 17, 20, 16 and 17 cells respectively, from at least 3 independent experiments). Scale bars, 2µm.

To enhance the formation and stability of cell contacts, we grew cells in linear microfabricated PDMS channels, optimized to constrain growth direction while providing large access to nutrients (Haupt, et al., 2018; Zegman, et al., 2015). As cells grew ends against ends in channels, they formed bright and long-lasting clusters at both old-ends and new-ends contact. These clusters were seen in 81% of ends contacts (n = 101 cells) and appeared more stable than in normal agar growth assay, presumably due to the absence of cell sliding, and/or the more crowded environment. In addition, clusters appeared more prominent at flat cell-cell interface, indicative of higher mechanical stress on the contact (Figure 1F-1G, Figure S1C-S1D and Movie S4). Further, by letting cells grow and divide in channels over longer periods of times, they adopted abnormal triangular or trapezoidal shapes, thereby pushing onto each other or onto the channel walls on lateral sides (Haupt, et al., 2018). Strikingly, Wsc1-GFP was systematically recruited to all lateral contacts, even those formed with channel walls, yielding clusters decorating a large part of cell sides (Figure 1H). In agreement with the formation of clusters against inert microchannels walls, Wsc1-GFP also clustered at cell-cell contacts with a *wsc1Δ* cell (Figure S1E-S1G). Thus, these results indicate that Wsc1 clustering may be triggered by local surface compression at any location around cells, independently of putative “*trans*” homotypic interactions between extracellular sensors from neighbor cells.

We quantified Wsc1 enrichment at cell contacts, by computing the average local signal to that of cell sides with no contact. Such quantification should yield a value of 2 for a contact-insensitive membrane factor, because of the presence of two apposed membranes at the contact site. Accordingly, the local enrichment of mCherry-Psy1, a single pass transmembrane SNARE, independent of any CW signaling, was 1.86 +/-0.58 (Figure S1H-S1J). These quantifications revealed a local enrichment ranging from ∼3 to 15-fold in Wsc1-GFP at sites of cell surface compression. Cluster sizes quantified as the full width at mid-height of a Gaussian fit of Wsc1 signal along the contour (Figure 1A), varied between ∼0.5 to 3µm in width, and were usually larger on larger contacts such as those formed along cell sides in microchannels (Figure 1I-1J). Importantly, larger clusters were not necessarily dimmer, suggesting that Wsc1 may form local sensing units that assemble into larger domains to decorate all the contact by further recruitment. Finally, this analysis revealed no clustering phenotype for the other surface sensor Mtl2, demonstrating it is specific to Wsc1 (Figure S1I-S1J).

As another situation implicating cell-cell contacts, we also imaged Wsc1-GFP dynamics during mating. Wsc1-GFP was generally enriched at mating tip projection, and became massively recruited when mating tips became apposed, reaching a local enrichment of up to ∼20 fold in a time-course of ∼20-80 min. Wsc1 clusters disappeared during cell fusion, as apposed CWs became digested (Figure 2A-2B and Movies S5) (Dudin, et al., 2015). Importantly, Wsc1 clusters still formed in mating pairs of *fus1Δ* cells, impaired in mating-specific polarized trafficking, but reached an enrichment, significantly smaller than in WT mating but similar to that found for interphase WT cell contacts (Dudin, et al., 2015). In addition, *gap1Δ* cells which form random mating projection without contacting partners did not enrich Wsc1 at cell tips (Figure 2A and 2C) (Merlini, et al., 2018; Imai, et al., 1991). Thus, Wsc1 clustering at mating tips may be driven by the contact of apposed CWs, and further enhanced by mating-specific polarity machineries. All together, these data demonstrate that the Wsc1 CW sensor may dynamically sense and accumulate to local cell-cell contacts where CWs are presumably pressed onto each other.

**Figure 2.**
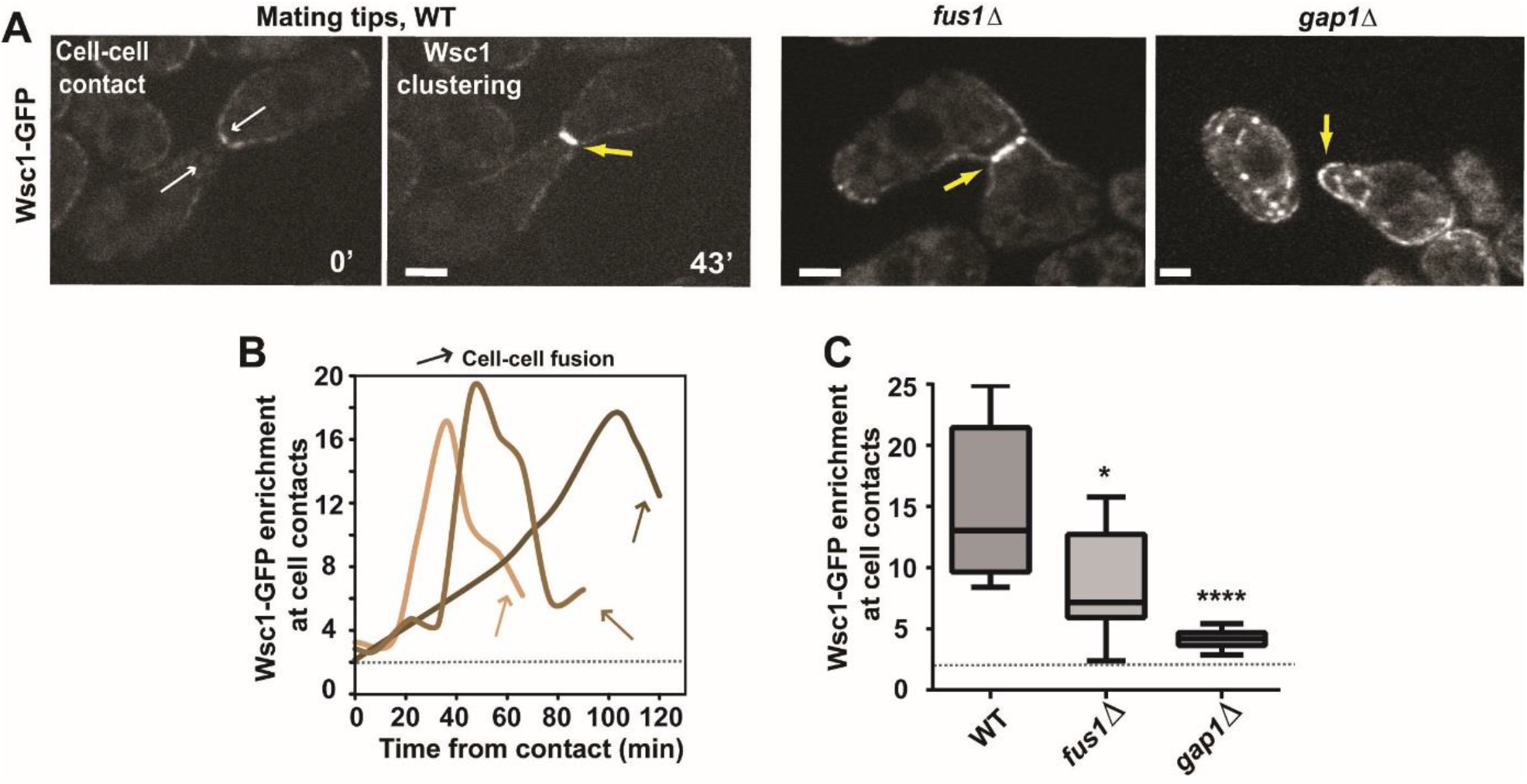
Wsc1 cluster formation at mating tips contacts. **(A)** Mid-slice spinning disk confocal time-lapse of WT cells expressing Wsc1-GFP undergoing mating (left). White arrows depict the direction of pressure-derived forces on the contact, and yellow arrows point to Wsc1 clusters. Mid-slice spinning disk confocal images of *fus1Δ* and *gap1Δ* mutants expressing Wsc1-GFP (right). **(B)** Dynamic of Wsc1-GFP enrichment in three representative pairs of WT mating cells. Time 0 corresponds to the first contact point, and arrows indicate the onset of CW opening prior to cell-cell fusion. The dotted line in the graph corresponds to a reference value of enrichment of 2, corresponding to the presence of two apposed membranes. **(C)** Quantification of Wsc1-GFP enrichment at cell contacts in the indicated conditions (n=13 cells for WT, 9 cells for *fus1Δ* mutant and 12 cells for *gap1Δ* mutant). For *gap1Δ* the enrichment was computed at shmoo tips and multiplied by 2 to compensate for the absence of partner cell. In all panels, yellow arrows point at Wsc1 clusters. Results were compared by using a two-tailed Mann–Whitney test. *, P < 0.05, ****, P < 0.0001, Scale bars, 2 μm.

### Wsc1 clustering is triggered by mechanical stress in the Cell Wall

Fission yeast cells harbor high internal turgor pressure on the order of ∼1-1.5MPa which impart large mechanical stress on the elastic CW (Abenza, et al., 2015; Atilgan, et al., 2015; Davi and Minc, 2015; Minc, et al., 2009). As cells grow and push onto each other, this stress may become enhanced locally, causing Wsc1 surface sensors to accumulate to cell contacts. To directly test the role of mechanical stress, we first followed Wsc1-GFP clusters formed in cells pressing onto each other in microchannels and rinsed with 2M sorbitol to drastically reduce turgor pressure. Cells rapidly deflated upon sorbitol treatment causing a complete disappearance of Wsc1 clusters within few minutes (Figure 3A-3B). Rinsing back with normal media, caused cells to re-inflate and Wsc1-GFP to re-accumulate at the same location over tens of minutes (Figure S2A-S2B). Because sorbitol treatment could indirectly affect Wsc1 membrane localization, we also followed Wsc1 clusters at cell contacts in microchannels, and ablated neighboring cells to relax mechanical stress at the contact (Haupt, et al., 2018). Following ablation, cells slowly separated, likely hindered by CW remnants of ablated cells or friction with channel walls, yielding to the progressive dissolution of Wsc1 clusters (Figure 3C-3D). Similarly, ablating one partner during mating, yielded cell shrinkage and CW relaxation in the non-ablated partner and cluster disappearance within few minutes (Figure S2C-S2D). Thus, these data directly suggest that Wsc1 forms reversible clusters at sites of enhanced mechanical stress in the CW, demonstrating that Wsc1 acts as a *bona fide* surface mechanosensor.

**Figure 3.**
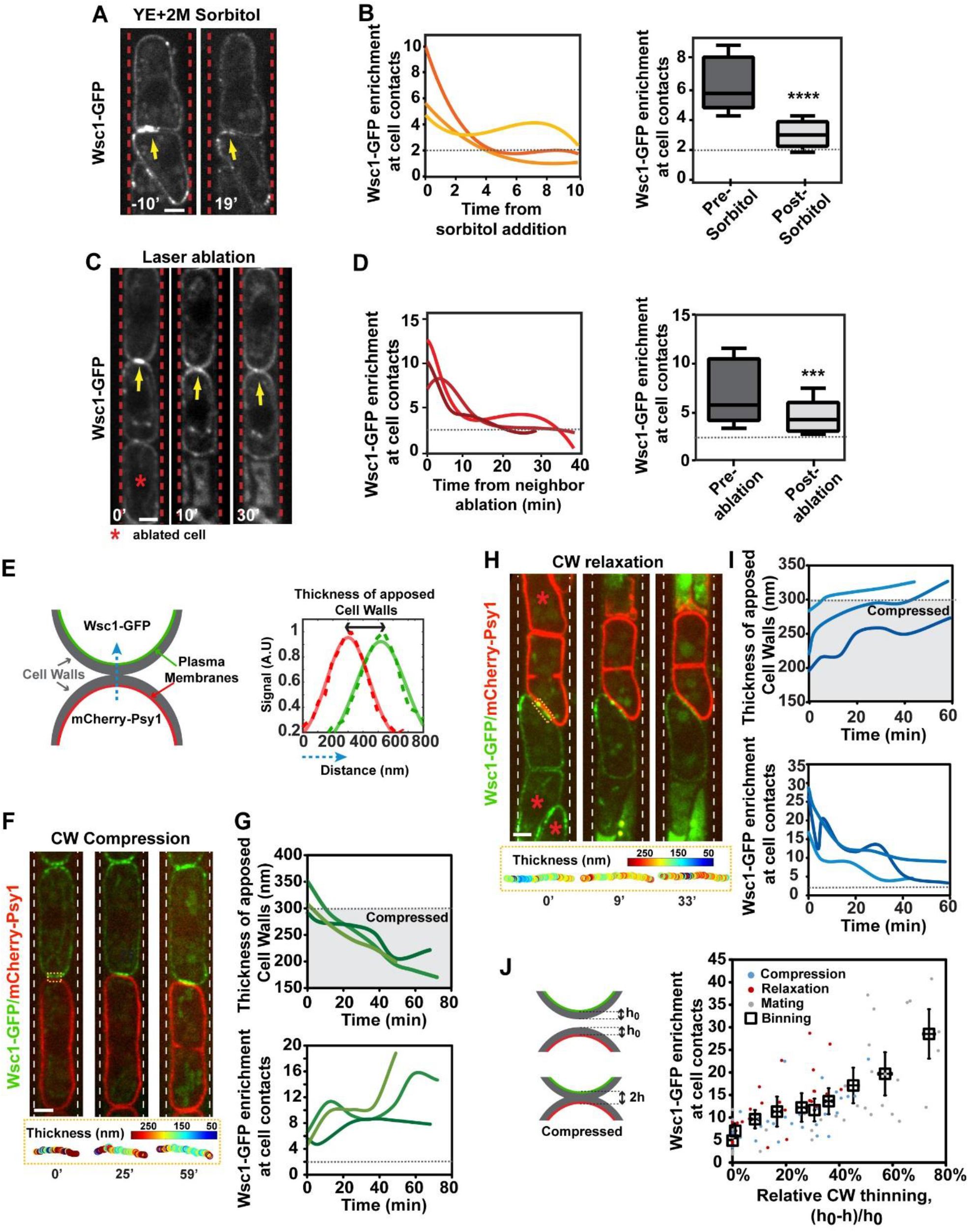
Compressive forces onto the Cell Wall trigger dose-dependence Wsc1 clustering. **(A)** Spinning disk confocal mid-section time-lapse of wild-type cells initially pressed onto each other through growth in microchannels and rinsed with 2M Sorbitol to reduce turgor pressure. Note the dissolution of Wsc1 clusters when cells deflate and relax stress on the cell wall. **(B)** Wsc1-GFP enrichment dynamics in three representative cells following sorbitol addition and box plots of Wsc1-GFP enrichment 10’ before and 10’ after sorbitol treatment (n= 5 cells). **(C)** Time-lapse of cells in microchannels, where laser ablation of one cell is used to relax pressure on neighbors, causing Wsc1 cluster dissolution. The red asterisk marks the ablated cell. **(D)** Wsc1-GFP enrichment dynamics following sorbitol addition and box plots of Wsc1-GFP enrichment 10’ before and 10’ after laser ablation of a neighbor (n= 5 cells). **(E)** Scheme representing the sub-resolution imaging method to compute the thickness of apposed cell walls from the peak signal of plasma membranes tagged with different fluorophores. **(F)** Example of a cell expressing Wsc1-GFP that grows into a cell expressing mCherry -Psy1 in microchannels, and computation of the thickness of CWs along the contact represented with a color map (bottom). **(G)** Thickness evolution of apposed cell walls as cells grow onto each other in three exemplary cell pairs, and concomitant evolution of Wsc1-GFP enrichment at the contact. The grey zone in the thickness plot corresponds to a zone smaller than the sum of normal thicknesses of CWs in free cells (Davì, et al., 2018). **(H)** Example of a cell expressing Wsc1-GFP pressed against a cell expressing mCherry-Psy1 in microchannels, where the contact is relaxed by ablating neighbors (marked with a red asterisk), and computation of the thickness of CWs along the contact represented with a color map (bottom). **(I)** Thickness evolution of apposed cell walls as cells separate and relax their cell walls at the contact due to neighbor ablation in three exemplary cell pairs, and concomitant evolution of Wsc1-GFP enrichment at the contact. **(J)** Wsc1-GFP enrichment plotted as a function of the relative thinning of apposed CWs with respect to free cells, for cells undergoing compression, relaxation in interphase or during mating. The squares are bins of 10 individual measurements, with error bars representing standard deviations (n= 92 clusters in total). Results were compared by using a two-tailed Mann–Whitney test. ***, P < 0.001, ****, P < 0.0001, Scale bars, 2 μm.

To directly assay the dose-dependent impact of CW mechanical compression on Wsc1 response, we adapted a previous sub-resolution method, to compute the sum of the thickness of apposed CWs at sites of cell contacts (Davi, et al., 2019; Davì, et al., 2018; Clark, et al., 2013). We mixed populations of cells expressing mCherry-Psy1 and cells expressing Wsc1-GFP in microchannels and monitored contacts at the interface between differently labeled cells. Because Wsc1-GFP is a single-pass transmembrane protein tagged at its intracellular tail, like mCherry-Psy1, we fitted each membrane signal with a Gaussian, and computed the peak-to-peak distance to extract the sum of thicknesses of apposed CWs (Figure 3E) (Davì, et al., 2018). Importantly our analysis script corrects for color spatial shifts and for variations in signal-to-noise, making the analysis mostly insensitive to the level of local signal enrichment (Davi, et al., 2019; Clark, et al., 2013). As controls, we verified that Wsc1-GFP signals gave similar values of mean CW thickness in normal cells, as GFP-psy1 (Figure S2E). In addition, measurements obtained with Wsc1-GFP or with a non-clustering membrane associated domain tagged with BFP (RitC-BFP), expressed in the same cell, yielded near similar apposed CW thickness values (Figure S2F-S2H) (Vjestica, et al., 2020). Using this method, we monitored concomitant changes in apposed CWs thickness and Wsc1 enrichments in time-lapses. As cells grew onto each other in channels, the contact flattened and the sum of apposed CWs transited from a value above ∼300 nm corresponding to the sum of normal CW thicknesses at cell tips (Davi, et al., 2019; Davì, et al., 2018), to values down to 150-200nm, corresponding to a relative thinning of CWs of up to ∼50%. This suggests that apposed CWs are submitted to compressive forces that cause them to thin with a rate likely determined by cell growth and divisions in the channel. Importantly, as CWs were more compressed, Wsc1-GFP clusters became more intense suggesting that progressive stress increase might promote further Wsc1 accumulation (Figure 3F-3G). Conversely, by performing neighbor ablation assays, CWs progressively relaxed yielding to the dissolution of Wsc1 clusters (Figure 3H-3I). Finally, the same analysis in mating cells, showed that apposed CWs of mating partners could thin by up to ∼80%, yielding more pronounced recruitment of Wsc1-GFP at CWs contacts (Figure S2I-S2J). During mating, this over-thinning is most likely partially accomplished by enzymatic digestion of apposed CWs by specific hydrolases in addition to pure mechanical thinning (Dudin, et al., 2015). Computing Wsc1-GFP enrichment as a function of CW thickness changes, for all above mentioned conditions, revealed a clear dose-dependence of Wsc1 enrichment as a function of CW compression (Figure 3J). Although the geometry of contacts and the presence of multiple cells precludes a simple calculation of the precise stresses borne by CWs, rough estimates based on the assumption that CW stress is inversely proportional to thickness (Davi, et al., 2019), suggest that an increase of ∼2X in local mechanical stress may drive a ∼10X enrichment in Wsc1-GFP. Thus, although these data do not presently allow discerning if Wsc1 directly senses thickness, elastic strains, stresses or other mechanical parameters, they strongly support a dose-dependent recruitment of Wsc1 sensors with compressive forces onto the CW.

### Wsc1 may behave as an autonomous module to detect local CW mechanical stress

Cytoskeletal elements, endo and/or exo-cytosis, as well as downstream signaling kinases, are recognized contributors of animal cell mechano-receptor clustering in response to force (Kechagia, et al., 2019; Changede and Sheetz, 2017; Welf, et al., 2012). We sought to dissect mechanisms of Wsc1 clustering upon mechanical stress. In normal growing cells, Wsc1-GFP localizes to cell tips, at sites largely enriched in polarity factors, actin assembly, endo- and exocytosis and CW remodeling regulators (Martin and Arkowitz, 2014; Cruz, et al., 2013). In addition, previous reports suggested that Wsc1 polar distribution is partially dependent on the actin cytoskeleton (Cruz, et al., 2013). Building on our microchannels assay where clusters are highly penetrant, and form at multiple sites around the cell surface, not only at cell tips, we first co-imaged active-Cdc42 (using the CRIB-GFP probe) as a general marker of polarity and actin assembly together with Wsc1-mCherry. When clusters formed at cell tips, they co-localized with CRIB-GFP, but were notably more focused than CRIB-GFP patches, accumulating at the precise contact where CW contacts appeared flat (Figure S3A). However, in a significant number of cells, clusters also formed at a contact with a neighbor CW away from the growing tip, creating a marked distinct localization between active-Cdc42 and Wsc1 cluster. Similar results were also obtained by co-imaging Wsc1 and the upstream regulator of polar growth and CW synthesis RFP-Bgs4 (Figure 4A). Thus, Wsc1 clusters forming at sites of local CW compression can be spatially uncoupled from polarity machineries.

**Figure 4.**
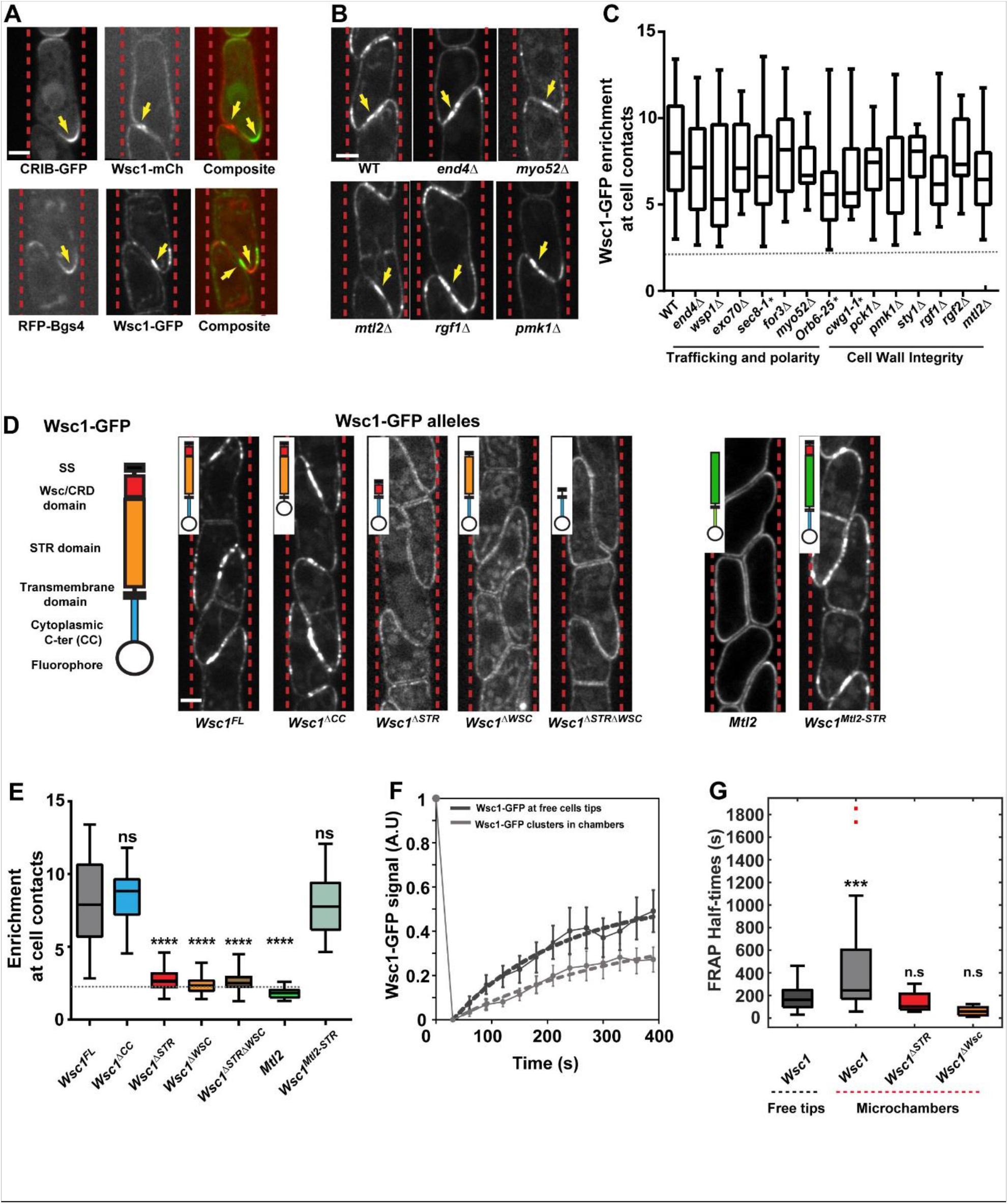
Force-dependent Wsc1 clustering is independent of downstream polarity, trafficking, and CW regulation, but requires extracellular CW interacting domains. **(A)** Co-imaging of active Cdc42 (CRIB-GFP) and Wsc1-mCherry clusters (top), and RFP-Bgs4 with Wsc1-GFP clusters (bottom) in cell pairs in microchannels. Note how Wsc1 clusters localize to sites of CW compression, while CRIB-GFP and RFP-bgs4 localize to growth sites. **(B)** Representative images of Wsc1-GFP clusters in cell pairs in microchannels in the indicated mutants. **(C)** Boxplots of Wsc1-GFP enrichment at sites of contacts in the indicated trafficking, polarity and cell wall integrity mutants and alleles (n > 35 cells for each condition, from at least two independent experiments). **(D)** Scheme representing the main functional Wsc1 domains, and representative spinning disk confocal mid-plane images of different tagged alleles lacking one or several of these domains, or with a domain swapped with the corresponding domain in Mtl2 (see insets) in microchannels. **(E)** Box plots of GFP-tagged alleles enrichment at cell contacts in microchannels. (*Wsc1* ^*FL*^*-GFP* (Full-length), n=65 cells; *Wsc1*^*ΔCC*^*-GFP* n= 45; *Wsc1*^*ΔSTR*^*-GFP* n= 66 cells; *Wsc1*^*ΔWSC*^ *-GFP* n= 33 cells; *Wsc1*^*ΔSTRΔWSC*^*-GFP* n= 87 cells, Mtl2-GFP n= 21 cells, *Wsc1*^*STR-Mtl2*^*-GFP* n= 26 cells). **(F)** Mean FRAP recovery curves of Wsc1-GFP at free tips or in clusters in microchannels (n= 28 and 27 cells, respectively, from 3 independent experiments). The dotted lines are exponential fits and error bars represent standard deviations. **(G)** Box plots of FRAP Half-times for the indicated conditions and alleles (n= 28, 27, 10 and 7 cells respectively). Red dots are outliers in the distribution. Results were compared by using a two-tailed Mann–Whitney test. n.s. P > 0.05, **, P < 0.01 ***, P < 0.001, ****, P < 0.0001, Scale bars, 2 μm.

We next screened a set of well-characterized mutants defective in endo- or exocytosis, actin assembly, secretory vesicle trafficking and polarity (Figure 4C and Figure S3B). Analyses of *end4Δ* and *wsp1Δ* mutants strongly defective in endocytosis, revealed no major defects or enhancement of Wsc1-GFP enrichment at compressed CWs. In addition, in these endocytic mutants, Wsc1 was largely delocalized from cell tips, suggesting that Wsc1 clustering does not require any prior tip localization. Similar clustering behaviors were obtained in secretion mutants, *exo70Δ* and *sec8-1*^*ts*^, as well as in the formin *for3Δ* and myosin type V *myo52Δ* mutants defective in secretory vesicle trafficking along actin cables (Martin and Arkowitz, 2014). This suggests that cluster formation may predominantly emerge as a result of reaction-diffusion or flows along the plasma membrane and CW, rather than recycling. An *orb6-25* ^*ts*^ mutant defective in polar growth, still formed clusters, confirming that Wsc1 does not rely on Cdc42-based polarity to form clusters at specific locations around cells. Together, these data suggest that Wsc1 senses and accumulates at sites of local CW compression independently of canonical polarity and trafficking pathways.

Because Wsc1 is upstream of the Cell Wall Integrity pathway (CWI), we also screened components of this pathway to test the contribution of putative positive feedbacks that could enrich Wsc1 clusters when the pathway is activated by surface forces (Figure 4C and Figure S3B) (Pérez, et al., 2018; Madrid, et al., 2006). Using a *cwg1-1* ^*ts*^ allele defective in CW synthesis, we first ruled out a major role of CW synthesis and remodeling in Wsc1 clustering. Also, null mutants in the protein kinase C, Pck1 and in the MAPK Pmk1 did not exhibit defects, suggesting no direct role for transcriptional responses. Similarly, upstream CWI GEFs Rgf1 and Rgf2 that activate the Rho GTPase Rho1 for CW repair did not exhibit notable defects in clustering. In addition, the other surface sensor Mtl2 did not contribute to Wsc1 clustering, further supporting their independence in probing and transducing CW stress (Cruz, et al., 2013). Finally, clusters also formed in a *sty1Δ* mutant defective in turgor adaptation, indicating that putative turgor pressure modulations may not contribute significantly to increased mechanical stress and Wsc1 clustering (Ryder, et al., 2019). Thus, downstream elements that regulate CW integrity and mechanical adaptation are not required for Wsc1 mechano-sensation.

### The CW-associated extracellular WSC domain is required for force sensing and clustering

To further dissect mechanisms and putative functions of Wsc1 clustering, we generated fluorescently tagged as well as untagged Wsc1 alleles that lack one or several of core functional domains: A Wsc1^ΔCC^ that lack the cytoplasmic C-terminal tail, a Wsc1^ΔSTR^ lacking the long STR “nano-spring” extracellular domain, a Wsc1^ΔWSC^ that lack the cysteine-rich WSC domain and a Wsc1^ΔSTRΔWSC^ that lack all extracellular domains except for the signal sequence. Allele design, followed sequence alignment with established alleles from *S. cerevisiae* Wsc1 (Philip and Levin, 2001), and all encoding constructs were integrated at the *wsc1*^+^ locus.

These alleles were viable, with no notable defects in cell size, division or morphology, but exhibited some low fraction (∼5%) of cell death similar to the null *wsc1Δ* mutant (see hereafter) (Cruz, et al., 2013). Importantly, tagged alleles behaved as untagged ones in terms of morphology and survival, and were still largely enriched at the plasma membrane, with Wsc1^ΔWSC^-GFP, exhibiting the most pronounced localization to internal membranes, potentially reflecting a role of the WSC domain for sensor stability at the membrane or within the CW (Kock, et al., 2015). Tagged-alleles were also all localized to cell tips, with Wsc1^ΔWSC^-GFP, Wsc1^ΔSTR^-GFP and Wsc1^ΔSTRΔWSC^-GFP exhibiting less polarized patterns, suggesting a partial minor function for extracellular domains in maintaining Wsc1 to the exact cell tips (Figure S3C-S3E). Using microchannel assays, we screened for putative clustering defects of these alleles. Remarkably, the Wsc1^ΔCC^-GFP lacking a large fraction of the C-ter, and thus presumably defective in downstream signal transduction was dispensable for clustering. This finding reinforces the notion that Wsc1 clustering occurs independently of downstream CWI signaling. In sharp contrast, Wsc1^ΔSTR^-GFP, Wsc1^ΔWSC^-GFP and Wsc1^ΔSTRΔWSC^-GFP were incapable of clustering, with an enrichment at contacts close to ∼2 similar to membrane proteins independent of CW signaling like Psy1 (Figure 4D-4E and Figure S1H-S1J). This indicates that WSC and STR domains are required for protein clustering under force. However, we note that given the long length of the rod-shape STR domains (estimated around 40-50nm, (Kock, et al., 2015; Dupres, et al., 2009)), one possible interpretation for the lack of Wsc1^ΔSTR^-GFP clustering, is that the WSC domain is placed at a much lower “altitude” in the CW, and cannot sense CW compression and properly promote clustering. To directly test this, we generated a Wsc1 allele in which the STR domain was swapped with that of the Mtl2 sensor, which does not form clusters. This swap construct exhibited a clustering phenotype similar to that of normal Wsc1 (Figure 4D-4E). We conclude that Wsc1 clustering under force is primarily driven by its extracellular WSC domain independently of cytoplasmic effectors.

### Wsc1 clustering through restricted diffusivity in the membrane and CW

To understand mechanisms driving clustering under force, we next performed Fluorescent Recovery After Photobelaching (FRAP) experiments. In cells growing in standard agar pads, Wsc1-GFP at free tips recovered with a half-time of ∼188.7 +/-117 seconds (mean +/-std), ∼1.2X higher than that of the integral membrane SNARE mCherry-Psy1 (Figure 4F and Figure S4A-S4C) (Bendezu, et al., 2015). FRAP images suggested that most of the recovery occurred along the membrane plane from lateral diffusive or advective transport (Figure S4B-S4C). Accordingly, FRAP half-times in an *end4Δ* mutant, defective in endocytosis, were nearly similar than in WT, suggesting that recycling does not contribute significantly to Wsc1 dynamics at these timescales (Figure S4A-S4C). Importantly, FRAP on clusters in microchannels revealed similar in plane motility, but yielded a half-time significantly higher than in free tips, of 457.3 +/-479 seconds (mean +/-std). In most clusters, we could observe clear lateral recovery and diffusion. In a subset, Wsc1 appeared almost frozen, with no detectable recovery within the cluster, and half-times exceeding 15-20 min timescales. We note that these half-times, are in the same order of magnitude, yet ∼2-3X faster than the typical timing for cluster formation seen in time-lapses when cells grow and press onto each other (Figure 1C,1E and 1G). We suggest that this difference reflects the slower force loading rate in these time-lapses, limited by cell growth and rearrangements. Finally, Wsc1^ΔWSC^-GFP and Wsc1^ΔSTR^-GFP incapable of clustering exhibited much smaller half-times closer to that of mCherry-Psy1 (Figure 4F-4G and Figure S4A). Together these results indicate that local mechanical stress on the CW may directly or indirectly hinder diffusion, or reaction-diffusion of Wsc1 sensors within membranes and CWs, potentially contributing to their clustering behavior.

### Functional relevance of Wsc1 clustering in cell survival

To assay the functional relevance of Wsc1 clusters, we next tested if they could recruit downstream elements of the CWI cascade. At free growing tips, Wsc1 co-localizes with the Rho1 GEFs Rgf1 and Rgf2, the active form of Rho1, and the protein kinases Pck1 and Pck2, all contributing to CWI pathway regulation (Pérez, et al., 2018; Garcia, et al., 2006) (Figure 5A). We co-imaged Wsc1-mCherry and Rgf1-GFP, ActRho-citrine, a marker for active GTP-bound Rho1, Pck1-GFP and Pck2-GFP (Figure 5A-5B). In multiple instances, we detected abnormal ectopic recruitment of these downstream factors at cell sides, in close association with a Wsc1-mCherry cluster formed at lateral CW contact with a neighbor or microchannel wall (Figure 5B). The incidence of these lateral zones was however relatively low (∼10-20% of Wsc1 lateral clusters), and in most cells these downstream elements were segregated to growing tips. We interpret this observation as a result of competition for CWI cytoplasmic effectors, with polar located ones that take over given their association to tip-located polarity cues (Pérez, et al., 2018). We conclude that Wsc1 clusters can serve as local “sensosome” platforms to recruit and activate downstream signaling elements to sites of mechano-perception.

**Figure 5.**
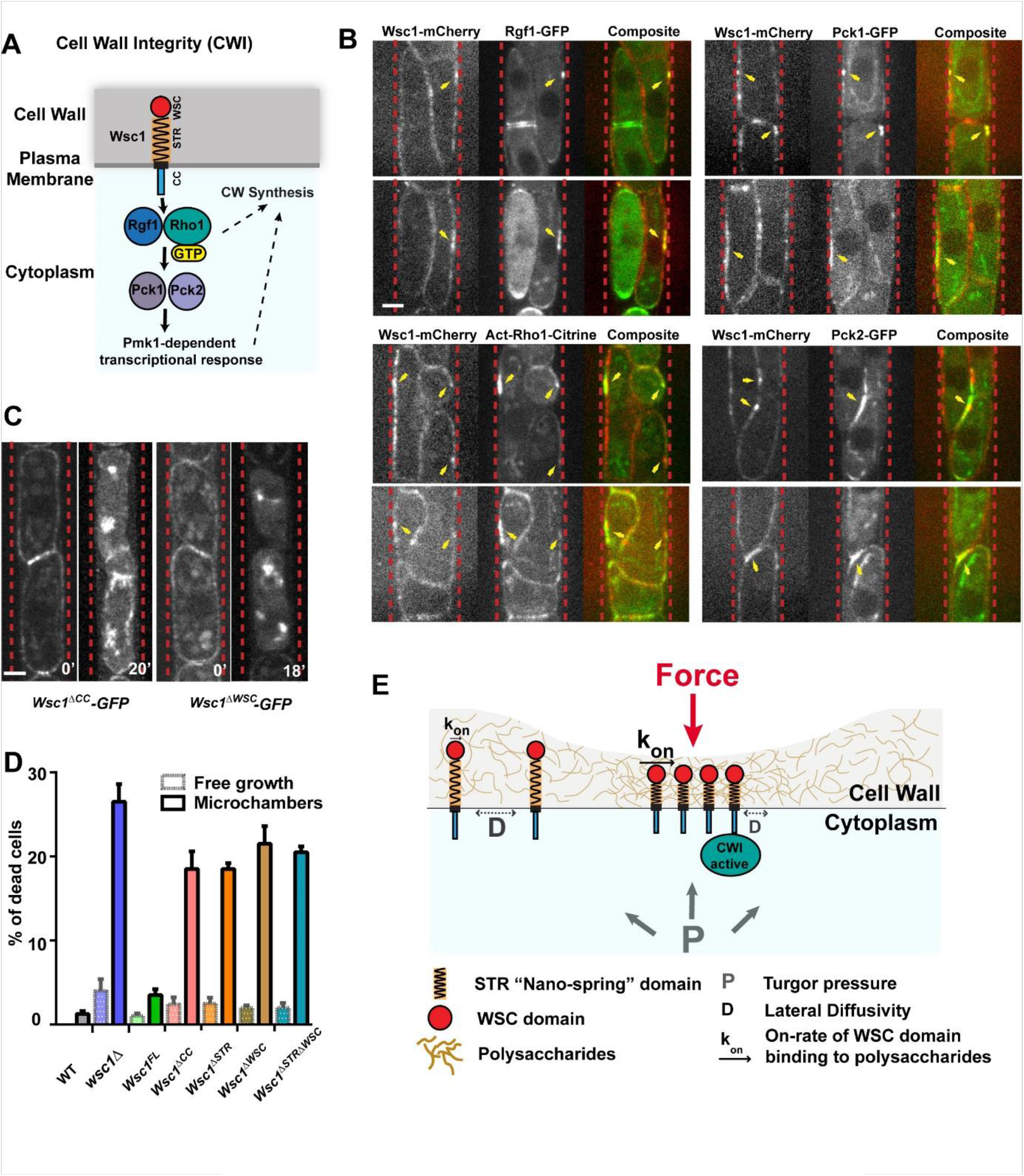
Force-dependent Wsc1 clusters can recruit downstream signaling factors and promote cell survival. **(A)** Scheme representing Wsc1 and its domains in the membrane and CW, and downstream cytoplasmic CWI components and regulation. **(B**) Representative spinning disk confocal mid-section images of cells grown in microchannels and co-expressing Wsc1-mCherry and representative upstream cytoplasmic effectors of the CWI (Rgf1-GFP, ActRho1-Citrine, Pck1-GFP and Pck2-GFP), featuring a clear visible ectopic accumulation of CWI components at lateral sites where Wsc1-mCherry form cluster due to CW compression (marked by yellow arrows) **(C)** Example of a pair of *Wsc1*^*ΔCC*^*-GFP* cells that form clusters at sites of enhanced surface stress, and undergo lysis. Example of a pair of *Wsc1*^*ΔWSC*^ *-GFP* cells which fail to form a cluster, at sites of enhanced surface stress and undergo lysis. **(D)** Quantification of cell death in different Wsc1 alleles, in normal agar pad (free growth) or in packed microchannels implicating higher values and frequency of CW mechanical stress (n ∼600 cells for each allele, from 2 independent experiments). **(E)** Proposed model for Wsc1 mechanosensation. Wsc1 diffuses in the membrane and CW planes, and interacts with CW polysaccharides through its WSC domain. Upon force application, the CW is locally compressed, promoting the densification of CW sugars. This may enhance the binding kinetics of WSC domains to sugars, slowing down lateral diffusivity, and leading to cluster formation. In parallel, at sites of CW compression, the STR domain becomes compressed and triggers downstream CWI activation through the cytoplasmic C-ter. Scale bars, 2 μm.

To test function of Wsc1 clusters more directly, we next assayed survival. In normal conditions, *wsc1Δ* exhibits a yield of ∼ 5-7% cell death by cell lysis (Cruz, et al., 2013). To determine which domain supports viability, we first computed cell death in all untagged versions of Wsc1 alleles. This revealed a yield of cell death of ∼4%, slightly lower than for *wsc1Δ* (Figure 5D). As Wsc1 is strictly required for survival in the absence of Mtl2 (Cruz, et al., 2013), we next used a shut-off construct for Mtl2, under the control of the low expression *81xnmt1* (No message in Thiamine) promoter. In plates supplemented with thiamine, to repress Mtl2 expression, *wsc1Δ, wsc1*^*ΔCC*^, *wsc1*^*ΔSTR*^ and *wsc1*^*ΔSTRΔWSC*^ were unable to grow. In contrast, *wsc1*^*ΔWSC*^ exhibited growth levels similar to wsc1 full-length, suggesting that the WSC domain is dispensable for survival in the absence of Mtl2, in normal growth conditions (Figure S5A-S5B). Thus, these results suggest that the *wsc1*^*ΔCC*^ is largely defective in downstream activation of the CWI, and that the STR “nano-spring” domain is needed for survival in the absence of the other sensor in normal growth conditions.

In microchannels, where clusters form at high frequency, due to larger and more frequent compressive mechanical stresses onto the CWs, the survival behavior was markedly different. First, *wsc1Δ* cells exhibited a much higher yield of death of ∼28%. Second, all alleles exhibited a high yield of death around ∼25%, including *wsc1*^*ΔWSC*^ which survives in normal growth conditions even in the absence of Mtl2. By imaging *wsc1*^*ΔCC*^-GFP, we could observe clusters that formed at a site of CW compression at tip-tip contacts, followed by death of the two apposed cells, presumably, because Wsc1 lacking its C-ter, was unable to properly recruit and activate the CWI pathway to strengthen the CW submitted to large mechanical stress. Similar death events could also be recorded in *wsc1*^*ΔWSC*^-GFP, where downstream signaling is likely intact, but given the absence of cluster formation (Figure 5C-5D). These results suggest that Wsc1 clustering is particularly relevant to growth in confinement, or in situations involving large numbers of cell-cell contacts and/or higher CW mechanical stress.

## DISCUSSION

### Mechanosensation in the cell wall

The mechanics of the CW is largely recognized to support survival, morphogenesis, infection and reproduction, in organisms ranging from bacteria to fungi and plants (Banavar, et al., 2018; Auer and Weibel, 2017; Davi and Minc, 2015; Ryder and Talbot, 2015; Boudaoud, 2010; Bastmeyer, et al., 2002). To date, however, the bare notion of mechanosensation in the CW remains underappreciated and understudied. In part, progress in this area has been limited by the lack of direct quantitative and genetically tractable readout of surface force sensing. By discovering that a conserved surface sensor embedded in the CW matrix, can re-localize and accumulate in massive clusters at sites of mechanical compression on the CW, our study thus constitutes an important step in the emerging field of walled cells mechanobiology. Clusters formed in a range of conditions, implicating surface forces, spanning tip growth, septation, mating and confinement, providing direct evidence, that even unicellular walled organisms can use surface mechanosensation to detect the presence of neighbors or obstacles. Importantly, our findings parallel with classical clustering phenotypes of mechano-receptors, such as integrins and cadherins, under force, which have been pivotal to dissect mechanisms of mechanosensation in animal cells (Kechagia, et al., 2019; Changede and Sheetz, 2017; Chen, et al., 2017; Truong Quang, et al., 2013; Galbraith, et al., 2002). Thus, we anticipate that our work could open the route to important studies on the mechanisms and functions of CW mechanosensation in yeast, fungi, and other walled cells.

### Surface sensors clustering

Seminal studies in *S. cerevisiae* had already reported on the formation of small Wsc1 foci/clusters, in normal growing cells, similar to what we observe in normal growing *S. pombe* cells (Figure S1) (Kock, et al., 2016; Heinisch, et al., 2010). Thus, the tendency of Wsc1 to form clusters along the cell surface, may be a built-in property of this protein and its interacting partners. However, our observation that these clusters can grow to micrometric sizes and form at specific sites where the CW is compressed, places Wsc1 as a *bona fide* mechanosensor, and raises the important question of how it may sense force. Our genetic analysis suggests that clustering under force, does not implicate canonical regulators of membrane trafficking, polarity and downstream CWI signaling. In addition, while removal of the Wsc1 cytoplasmic C-ter intracellular tail does not affect clustering under force, removal of any of the CW-associated extracellular domains completely abolishes it. In addition, swapped STR domain with non-clustering Mtl2 analysis, suggest that the WSC domain is the main extracellular domain to promote clustering. These data strongly suggest that most of the ability to sense and accumulate in response to forces is solely encoded in the extracellular Wsc1 domains, placing Wsc1 as a potential autonomous sensor of surface forces.

One previously proposed mechanism for Wsc1 clustering in *S. cerevisiae*, is that the high content of Cysteine residues in the WSC domain may favor protein-protein cis-interactions through disulfide bond formation (Kock, et al., 2016). However, our observation that clusters can dissolve within few minutes when CWs are relaxed, do not simply align with the role of covalent interactions in cluster formation under force. Rather, our FRAP data and allele analysis supports a model based on restricted local diffusivity, and weak interactions between the WSC domain and CW polysaccharides. In that view, we posit that local CW compression could modulate the density/arrangement of the CW matrix favoring the binding of WSC domains to polysaccharides, thereby reducing lateral diffusivity and favoring clustering (Figure 5E). In addition, as the STR “nano-spring” domain mediates downstream transduction, we propose that its natural conformation is stretched, and that it compresses along with the CW, when force is applied, presumably driving conformational changes and/or phosphorylation of the cytoplasmic C-ter for CWI activation (Figure 5E). Accordingly, AFM-based measurement stiffness of the STR suggests it is 3-4 orders of magnitude softer than the CW, so that its length should directly follow the position of the WSC domain above that moves transversally in response to force. Our proposed model has conceptual equivalence to “kinetic trap” models for integrin clustering, in which integrin interactions with ECM ligands reduces lateral diffusivity and promote clustering (Kechagia, et al., 2019; Welf, et al., 2012; Paszek, et al., 2009; Ballestrem, et al., 2001; Laukaitis, et al., 2001). Importantly, our model could presumably allow Wsc1 to detect multiple CW parameters, including thickness, elasticity, stress, or strain, which cannot be easily tested in our experiments. We anticipate that finer molecular dissection of extracellular domains, local manipulation of CW matrices, as well as more resolved methods to control force application onto the cell surface, may help to test and expand these initial CW mechanosensing models.

### Implication for cell physiology and survival

Clustering is a conserved feature of surface and intracellular sensors to form “sensosome” platforms, which serve as hubs for signal perception and transmission. Accordingly, our data suggest Wsc1 clusters can recruit downstream signaling elements at sites of mechanical compression and promote survival under large and more frequent stresses encountered in microchannel assays. We note that we could not assay transcriptional activation given that *S. pombe* Pmk1 localizes to both cytoplasm and nucleus independently of stress-induced activation of the pathway (Madrid, et al., 2006). The physiological relevance of microchannels to natural yeast or fungal habitat may include dense colonies or biofilms. In addition, we provide significant evidence that clusters form even in classical dilute growth assays, albeit more transiently, at sites of cell-cell contacts, when cells grow and collide onto each other’s at the end of septation, or at growing tips in interphase. In addition, smaller dynamic clusters also appear even at free growing tips, independently of any contact. Although we have not yet explored the significance of these clusters, we suspect that they could correspond to local sites of CW thinning or higher stress (Figure S1A). Wsc1-mediated mechanosensation could thus represent a basal homeostatic module contributing to homogenize CW mechanics in time and space at remodeling growth sites (Banavar, et al., 2018; Davì, et al., 2018). Finally, we also observe massive clustering at sites of mating tip contacts. MID-type channels were first identified from the analysis of “Mating Induced Death” mutants in *S. cerevisiae* (Ono, et al., 1994), and functions of WSC-types sensors have also been recently suggested for mating in these cells (Banavar, et al., 2018). The accumulation of Wsc1 at mating tip contacts, in *S. pombe*, suggests it could act to slow down or buffer CW digestion. Future studies incorporating the notion of CW mechanosensation in the regulation of growth, reproduction, survival, or microbial infection promise to highlight the role of mechanical feedbacks in the basic lifestyles of walled cells.

## ACKNOWLEDGMENTS

We thank Sophie Martin, Pilar Perez and Kathy Gould’s labs for sharing strains and reagents, as well as our colleagues Arezki Boudaoud and Sebastian Léon for carefully reading this MS. We gratefully thank all members of the Minc and Sanchez teams for discussion and technical help. This work was supported by the MEIC, Spain (BFU2017-84508-P) and the Regional Government of Castile and Leon [SA073U14] to Y. S, and the Centre National de la Recherche Scientifique (CNRS), the Agence Nationale pour la Recherche (“Cell size” no. ANR-14-CE11-0009-02 and “CellWallSense” no. ANR-20-CE13-0003-01) and the European Research Council (ERC CoG “Forcaster” no. 647073) to N.M.

## AUTHOR CONTRIBUTIONS

Conceptualization, N.M., Y.S. and R.N.V.; Methodology, N.M., Y.S., R.C. and R.N.V. Writing –Original Draft, N.M., and R.N.V. Draft Editing. N.M., Y.S., R.C. and R.N.V

## MATERIAL AND METHODS

### Fission yeast strains, growth conditions and genetics

Standard *S. pombe* genetic procedures and media were used (Forsburg and Rhind, 2006; Moreno, et al., 1991). The relevant genotypes and the source of the strains used are listed in Supplementary Table 1. Genetic crosses and selection of the characters of interest was done by random spore analysis. All the deletions strains generated in this work were checked by PCR, drop assays with caspofungin (0,5-1 µg/ml) and the appropriate selection markers. For all microscopy experiments, unless otherwise indicated, liquid cultures were grown overnight in YE5S at 25°C, diluted in fresh medium and grown to an optical density (OD_600_) between 0.4 and 0.6 before live-imaging.

i. Wsc1-mCherry was obtained by tagging at its endogenous genomic locus at the 3′ end, yielding a C-terminally tagged protein. This was achieved by PCR amplification of a fragment from the template plasmid with primers carrying 5′ extensions corresponding to the last 80 nucleotides of the Wsc1 ORF and the first 80 nucleotides of the 3′UTR and integrated in the genome by homologous recombination, as previously described (Bahler, et al., 1998). The plasmid pVD11 (pFA6a-*mCherry*-*kanMX*) was used as a template for PCR with primers RN01 (5′-<u>tgatgaatcgcaaatcaagtgaatctttggctgacagtcaggattattcgagaaagatattgcgtgtcacaaatttgaac</u>cggatccccgg gttaattaa) and RN02 (5′-<u>attatgaaagtcataaacaccataaagaagaaataattttttggttgaatttaaataaagtaaaaaagaaaaatgttgtt</u>gaattcgagctcgt ttaaac) (*wsc1* homology underlined) and the resulting PCR product was transformed into a Wild-type (WT) h90 strain. The positive clones were selected on YE5S + 120 µg/mL Kan and the integration was confirmed by PCR.
ii. Marker switching from Kan to Nat was achieved by amplifying the NatMX cassette from a pCR2.1-nat vector with primers MD1 and MD2 and transforming it into KanMX cassette-bearing strains, as described in (Haupt, et al., 2018).
iii. To generate *fus1Δ* and *gap1Δ* mutants expressing Wsc1-GFP, we used the plasmids pSM1966 (pFA6a-*fus1*-3’UTR-AfeI-5’UTR-*kanMX*) and pSM1869 (pFA6a-*gap1*-3’UTR-AfeI-5’UTR-*hphMX*) (kind gift from S. Martin, University of Lausanne, Switzerland). These plasmids were linearized with the restriction enzyme AfeI, purified, and transformed into a WT homothallic strain (RN42, h90). Positive clones were selected on YE5S + 120 µg/mL Kan (or) YE5S + 300 µg/mL Hygromycin B plates and tested for functional and visual phenotypes.
iv. Generation of Wsc1 alleles. The mutated versions of Wsc1 were based on pAL-*wsc1*^+^-GFP (pRZ21, wsc1 GFP-tagged in a *Not*I site at the C-end with the *ura4*^+^ marker inserted at the 3énd (Cruz, et al., 2013)). To create *Wsc1*^*ΔWSC*^ (pRC22), pRZ21 was modified by site-directed mutagenesis with a deletion loop oligonucleotide that eliminates base pairs (bp) from positions +93 to +351 (5’-GGTTTGCAAAACCCCATTGCCAGTAGCGCTCATGTCAGCCGCCACCAACCGAGT-3’). For *Wsc1*^*ΔSTR*^ (pRC23) we eliminated base pairs from positions +354 to +846 with oligonucleotide (5’-GCATTCAAGGACGTATGGTTTGAAGTCAGATAAACCGACCAGTACAG-3’) and for *Wsc1*^*ΔSTRΔWSC*^ (pRC24) a fragment (+93 to +846) was eliminated with oligonucleotide (5’-GCATTCAAGGACGTATGGTTTGACATGTCAGCCGCCACCAACCGAGT-3’). Finally, *Wsc1*^*ΔCC*^ (pRC25) was obtained with a deletion loop oligonucleotide that eliminates base pairs from positions +969 to +1098 (5’-CAAATTTGTGACACGCAATATTCTAATTTTGAATCTCCTGAA-3’). From each plasmid, an *Spe*I-*Bgl*II fragment containing the mutagenized *wsc1* open reading frame (ORF) and flanking sequences was transformed into the wild type strain (SM215). Thus, each construct was present as a single copy in the cell under the control of native Wsc1 promoter. To make the untagged mutated versions of Wsc1, plasmids pRC22, pRC23, pRC24 and pRC25 were closed with *Not*I to eliminate the GFP epitope. The genomic mutated alleles were tested by PCR to validate if they hold the right deletion and if they replace the wild-type allele. To obtain the *wsc1* mutants (*wsc1* ^*ΔWSC*^, *wsc1* ^*ΔSTR*^, *wsc1* ^*ΔWSC ΔSTR*^ and *wsc1* ^*ΔCC*^) in the *mtl2*^+^ shut-off background, a cassette (*Spe*I-*Bgl*II fragment) for each *wsc1* deletion allele (untagged) was used to transform SC80 (*kan-P81xnmt1-mtl2*^*+*^) cells carrying the *mtl2*^+^ gene driven by the P81x*nmt1* promoter (Cruz, et al., 2013).
v. Generation of Mtl2-Wsc1 STR swap construct GFP-Wsc1CC-Wsc1TMD-Mtl2STR-Wsc1CRD-Wsc1SS was based on pRZ21 (*wsc1*^*+*^). pRZ21 was modified by site-directed mutagenesis by introducing two *Sal*I sites at positions +351 and +846 bp to eliminate the STR domain. To isolate the STR fragment from *mtl2*^+^, pSC19 (Cruz, et al., 2013) was mutagenize to create two *Sal*I sites at positions +105 and +702. Then the STR fragment from *mtl2*^+^ (flanked by *Sal*I sites) was inserted in frame into pRZ21 cut with *Sal*I. As above, an *Spe*I-*Bgl*II fragment containing the hybrid *wsc1-STR-mtl2* was used to transform the wild type strain (SM215).

### Mating assays

Mating assays were performed as described in (Vjestica, et al., 2016). Briefly, liquid cultures of homothallic strains were first grown in a minimum sporulation medium supplemented with Nitrogen (MSL+N) at 25°C until reaching an optical density (OD_600_) between 0.5 and 1.0. On the second day, cultures were then diluted to an OD_600_∼ 0.025 in MSL+N and grown for 16-20h at 30°C to reach an OD_600_∼ 0.5-1.0. The third day, cells were washed thrice with MSL-N (without Nitrogen), and re-suspended to an OD_600_∼ 1.5 in 1 ml MSL-N and incubated at 30°C for 4-6h under agitation. Next, the cells were pelleted down by centrifugation, and 1 µl of cells were directly placed between an MSL-N agarose pads and a coverslip, followed by sealing with VALAP. The slide containing the cells were incubated for ∼15h in a humidified petri dish at 18°C. On the next day, microscopy images at different stages of mating were captured. For mating experiments between partners expressing different markers of the membrane, needed to compute apposed thickness of cell walls at sites of mating tip contact, heterothallic strains of Wsc1-GFP (h-) and mCherry-Psy1 (h+) were grown in separate tubes as described above until the second day. On the third day an equal number of heterothallic partners were mixed, centrifuged and re-suspended in MSL-N medium. The next subsequent steps were similar as above.

### Micro-fabricated channel handling

PDMS microchannels were fabricated and mounted as described in (Haupt, et al., 2018). For experiments using microchannels, cells were loaded as spores, which then germinated and proliferated as vegetative cells and crowded the microchannels (Haupt et al., 2018). Spores were obtained from homothallic h90 strains after placing freshly growing cells on solid ME medium plates at 25°C for 2-3 days. Mating mixtures were digested in 1/200 Glusulase (PerkinElmer, Waltham, Massachusetts, USA) at room temperature overnight, to eliminate vegetative cells. Digests were cleared of debris by adding four volumes of Percoll (Sigma-Aldrich) followed by centrifugation. Fresh spores were re-suspended in YE5S and pushed into channels by applying a pressure on the channel inlet with a syringe, usually yielding 1-6 spores per channel. The spore-containing channels were filled with YE5S and incubated for 16-18h in a humidified petri dish at 25°C. The next morning, channels were rinsed with fresh YE5S at least 1h before live-imaging. For temperature sensitive alleles, *cwg1-1, orb6-25*, and *sec8-1* spores were grown in channels at permissive temperature (25°C) overnight, rinsed with fresh media the next morning and then switched to restrictive temperature (36°C) for 6h, 2h, and 1h respectively, before observation. For sorbitol treatments, cells were grown in microchannels in YE5S medium as above, and subsequently rinsed with large volumes of YE5S + 2M Sorbitol, to ensure fast medium exchange. To compute the thickness of apposed cell walls at sites of cell contacts in microchannels an equal volume of Wsc1-GFP and mCherry-Psy1 expressing spores were mixed and pushed into the channels. These channels were filled with YE5S and incubated for 16-18h at 25°C.

### Cell death assays

For cell death assay experiments using untagged Wsc1 alleles, homothallic strains were either grown in microchannels or in liquid cultures in YE5S medium. Cells grown in liquid cultures, were placed on an agarose pad, and imaged within 1h, and cell death was quantified visually. For microchannel assays, cells were grown to a high confinement, assayed from the appearance of mis-shaped cells, and cell death was quantified visually. For cell death assay experiments using untagged Wsc1 alleles in an Mtl2*-*shutoff background under the control of the P81x*nmt1* promoter, liquid cultures were grown overnight in Edinburgh Minimal Medium supplemented with amino acids at 25°C. The next morning, 2 ml of this overnight culture was pelleted and diluted in Edinburgh Minimal Medium supplemented with amino acids and with 5 μg/ml of thiamine with a starting OD_600_ ∼ 0.05-0.1. The cells were left in the incubator at 25°C for ∼9h before imaging.

### Microscopy

All live-cell imaging experiments were performed on YE5S 2%-agarose pads, or MSL-N 2% agarose pads (for mating) or in microchannels. Imaging was carried out at room temperature (22–25°C) on an inverted spinning-disk confocal microscope equipped with a motorized stage and an automatic focus (Ti-Eclipse, Nikon, Japan), a Yokogawa CSUX1FW spinning unit, and an EM-CCD camera (ImagEM-1K, Hamamatsu Photonics, Japan). Images were acquired with a 100X oil-immersion objective (CFI Plan Apo DM 100X/1.4 NA, Nikon) in combination with a 2.5X magnifying lens. For laser ablation assays, we used an iLas2 module (GATACA Systems, France) in the ‘‘Mosquito’’ mode, irradiating cells at diffraction-limited spot with a 355 nm laser with a 60X oil-immersion objective (CFI Apochromat 60X Oil λS, 1.4 NA, Nikon), and the subsequent images were captured using a 100X oil-immersion objective. The microscope was operated with Metamorph (Molecular Devices). All images and time-lapses presented are single confocal mid-slice.

Fluorescence Recovery after Photobleaching (FRAP) was performed using the iLas2 module (GATACA Systems, France) on the spinning disk confocal system described above. A 1.0 μm circular ROI was bleached following two pre-bleach acquisitions and recovery was followed for every 30 second interval. Other shorter or longer time-intervals were also assayed to optimize this analysis and identify the proper timescale to quantify Wsc1 motility (Data not shown).

Cell death assays were performed on a wide-field fluorescence microscope (TI-Eclipse; Nikon) operated with Micro-Manager (Open Imaging). Cells were imaged with a 100X objective in DIC (Differential Interface Contrast) which allows to clearly distinguish dead from live cells.

### Image analysis and quantifications

To quantify local fluorescent enrichment, a semi-automated home-made Matlab (Mathworks) script was developed. The script removes fluorescent background and allows the user to trace a line manually around the cell contour, visible from Wsc1-GFP or other marker’s signals, around cell tips, or cell sides. The fluorescence was averaged on a thickness of 14px across the membrane (for a pixel size of 43.1nm), in order to average signals from both cells at sites of cell contacts, and fitted with a Gaussian, to extract basal fluorescent levels away from clusters, and maximum signal in clusters. The ratio between max and min, was used to compute enrichment and the full width at mid-height to calculate cluster sizes. In order to compare Wsc1 enrichment, at a contact between two cells, with that in a free cell tips, or to a contact formed with a neighbor expressing a protein tagged in another color, the scripts was modified in the latter cases to a thickness contour of 7 px, and the enrichment was multiplied by a factor 2.

FRAP data were analyzed, by computing the fluorescence recovery in the bleached region, after background subtraction, and was normalized to the signal of 3 unbleached regions of interest in cells in the same field, to account for photobleaching. Each recovery curve was fitted with a single exponential, to extract individual half-times and mobile fractions. The data were also averaged to plot mean recovery dynamics presented in Figure 4.

Cell wall thickness measurements were performed as described in (Davi, et al., 2019; Davì, et al., 2018). Briefly this analysis initiates with a two-color mid-slice confocal image, and traces segments perpendicular to the membrane, to detect the local distance between the two Gaussian peaks. Upon image registration, obtained by calibrating the field with multi-spectral beads scanned around the field of view, the distance is corrected and computed to extract local thickness. The impact of differences in signal intensities are automatically corrected to extract the true signal peak by using an analytical expression of the convoluted intensity profile. Thicknesses of apposed cell walls were averaged around the center of the zone of cell contacts where clusters formed.

### Statistical analysis

All experiments presented in this MS were repeated at least twice and quantified in a number of cells or events detailed in each figure legend. Statistical and correlation analyses were carried out using Prism 6 (GraphPad Software, La Jolla, CA). To compute significance throughout this work, we used, one-way analysis of variance (ANOVA) followed by Dunnett multiple comparisons test or two-tailed, unpaired t test. Statistically significant differences are reported in figure legends.

## SUPPLEMENTAL MATERIAL

Supplemental material contains 5 Supplemental Figures and Figure Legends, Supplemental Movie legends and 1 Supplemental Table.

**Figure S1(Related to Figure 1).**
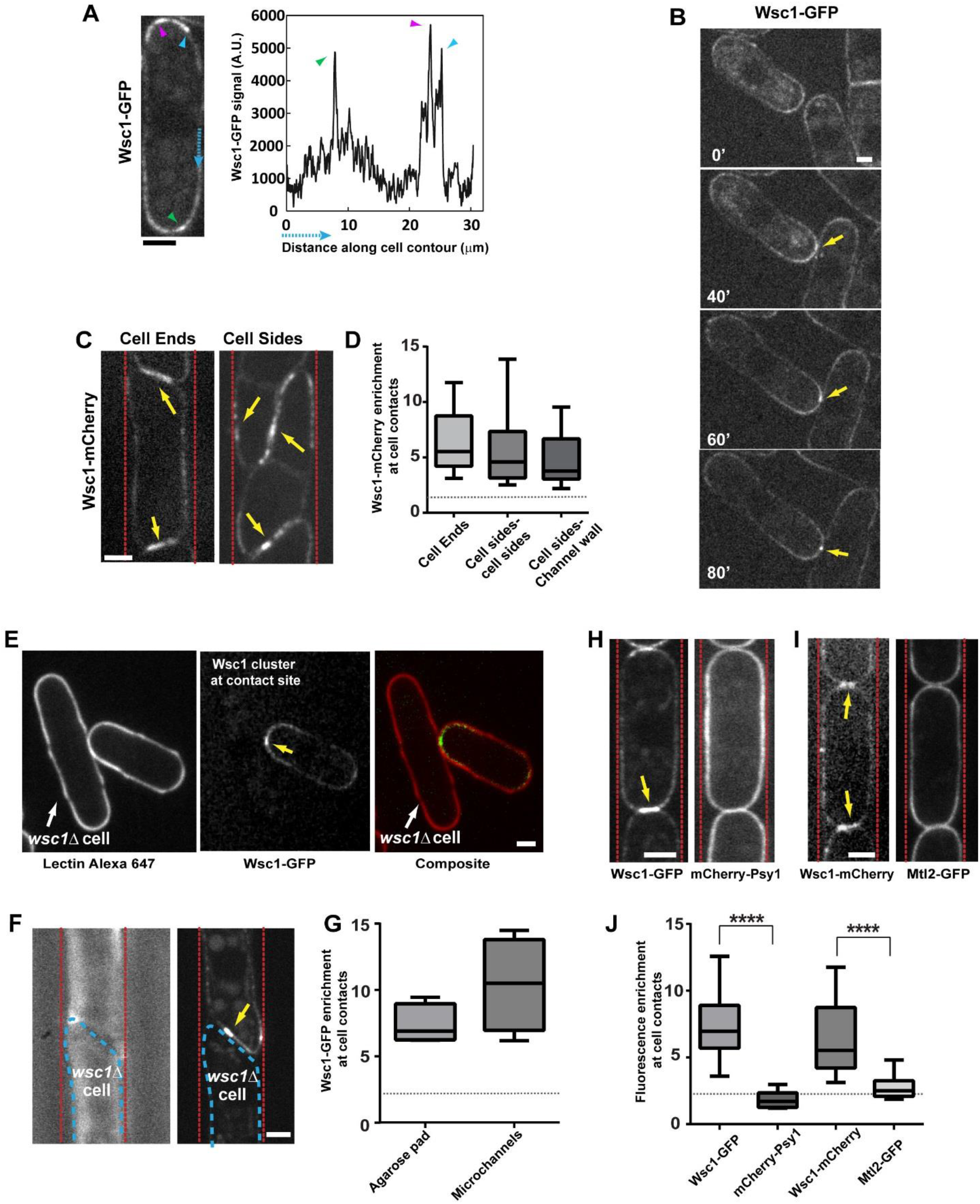
Cluster formation at cell contacts is specific to Wsc1 and independent of putative “trans” interactions between sensors from adjacent cells. **(A)** Mid-slice confocal image of a WT cell expressing Wsc1-GFP, with colored arrowheads pointing at local foci enriched in fluorescence, and corresponding fluorescent signal traced around the cell contour. **(B)** Time-lapse of WT cells growing onto each other, forming a cluster at the site of contact between cell tip and cell side, that dynamically tracks the site of contact. **(C)** Clusters formed by Wsc1-mCherry at cell ends contacts, and lateral contacts with other cells or with channel walls in microchannels. **(D)** Quantification of Wsc1-mCherry fluorescent enrichment at cell contacts (n=16 cells for cell ends and cell sides-cell sides, and 12 cells for cell sides-channel wall). The dotted line marks a reference enrichment of 2. **(E-F)** Wsc1-GFP cluster at a contact with a *wsc1Δ* cell in agarose pads (E), or in microchannels (F). **(G)** Fluorescence enrichment, multiplied by 2, to normalize for the absence of signal in the neighbor cell (n=5 cells for the two conditions). **(H)** WT cells co-expressing Wsc1-GFP and mCherry-Psy1 (an integral membrane protein independent of cell wall regulation), forming contacts in microchannels. **(I)** WT cells co-expressing Wsc1-mCherry and Mtl2-GFP (the other surface sensor of the cell wall integrity pathway) forming contacts in microchannels. **(J)** Quantification of fluorescent enrichment for the indicated markers at cell contacts in microchannels (n=15 cells for Wsc1-GFP /mCherry-Psy1, n=9 cells for Wsc1-mCherry/Mtl2-GFP). The dotted line marks a reference enrichment of 2. In all panels, yellow arrows point at Wsc1 clusters. Results were compared by using a two-tailed Mann–Whitney test. ***, P < 0.001, ****, P < 0.0001, Scale bars, 2 μm.

**Figure S2(Related to Figure 3).**
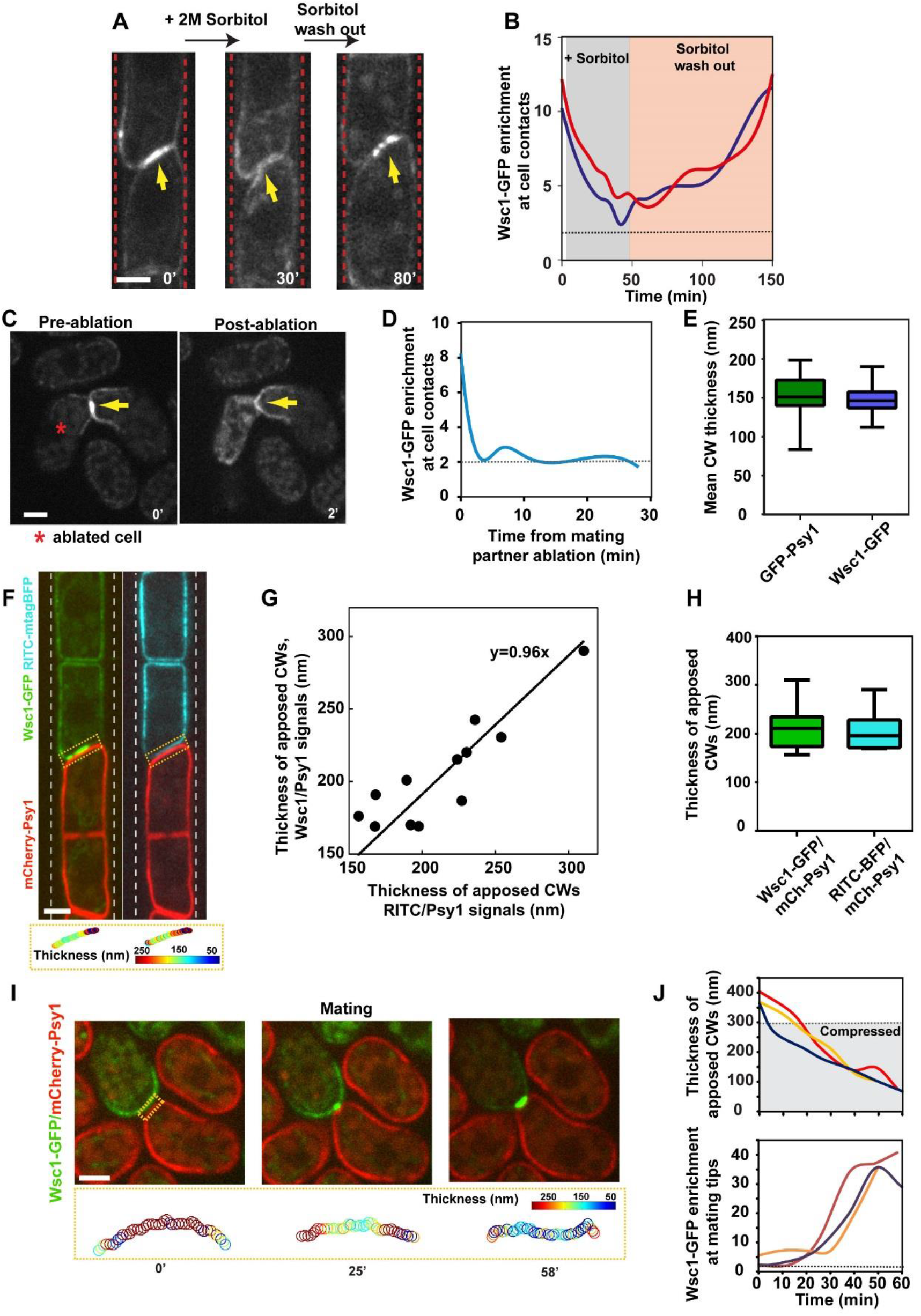
Wsc1 clusters in response to surface forces onto the Cell Wall. **(A)** Mid-slice spinning disk confocal time-lapse of a pair of WT cells with a Wsc1-GFP cluster in microchamber, rinsed with 2M sorbitol to reduce turgor pressure, and subsequently rinsed back in YE5S medium, to recover turgor pressure. **(B)** Dynamic of Wsc1-GFP enrichment in two representative pairs of WT cells in microchannels, rinsed with 2M sorbitol and then with YE5S medium. The dotted line in the graph corresponds to a reference value of enrichment of 2, corresponding to the presence of two apposed membranes. **(C)** Mating WT cells, in which one of the partner was ablated with a laser, to release stress on the CW at the site of cell contact. **(D)** Dynamic of Wsc1 enrichment at mating tip contacts in the experiment presented in (C). **(E**) Quantification of mean cell wall thickness around WT cells using sub-resolution methods as in (Davì, et al., 2018) measuring the distance from GFP-psy1 or Wsc1-GFP to a fluorescent lectin Gs-IB4-Alexafluor647 (n=25 cells for each condition). **(F)** Mid-slice spinning disk confocal images of a pair of WT cells formed between one cell expressing Wsc1-GFP and RitC-BFP and a cell expressing mCherry-psy1, and colored map of computed cell wall thickness at the contact site. **(G)** Thickness of apposed cell walls at contact sites in microchannels, measured from the sub-resolved distance between Wsc1-GFP and mCherry-psy1 signal peaks, plotted as a function of the same thickness measured from the distance of Ritc-BFP and mCherry-psy1. Each point represents one contact between a pair of cells as in F. The line is a linear fit, with the slope indicated in the graph (n=12 cell pairs). **(H**) Box plot of data presented in G. **(I)** Time-lapse sequence of mating between a cell expressing Wsc1-GFP and a cell expressing mCherry-psy1, and thickness of apposed cell walls along the contact at the bottom. **(J)** Thickness evolution of apposed cell walls in mating pairs that contact and press onto each other in three exemplary cell pairs, and concomitant evolution of Wsc1-GFP enrichment at the contact. The grey zone in the thickness plot corresponds to a zone smaller than the sum of CW thicknesses in free mating tips (Davì, et al., 2018). In all panels, yellow arrows point at Wsc1 clusters. Scale bars, 2 μm.

**Figure S3(Related to Figure 4).**
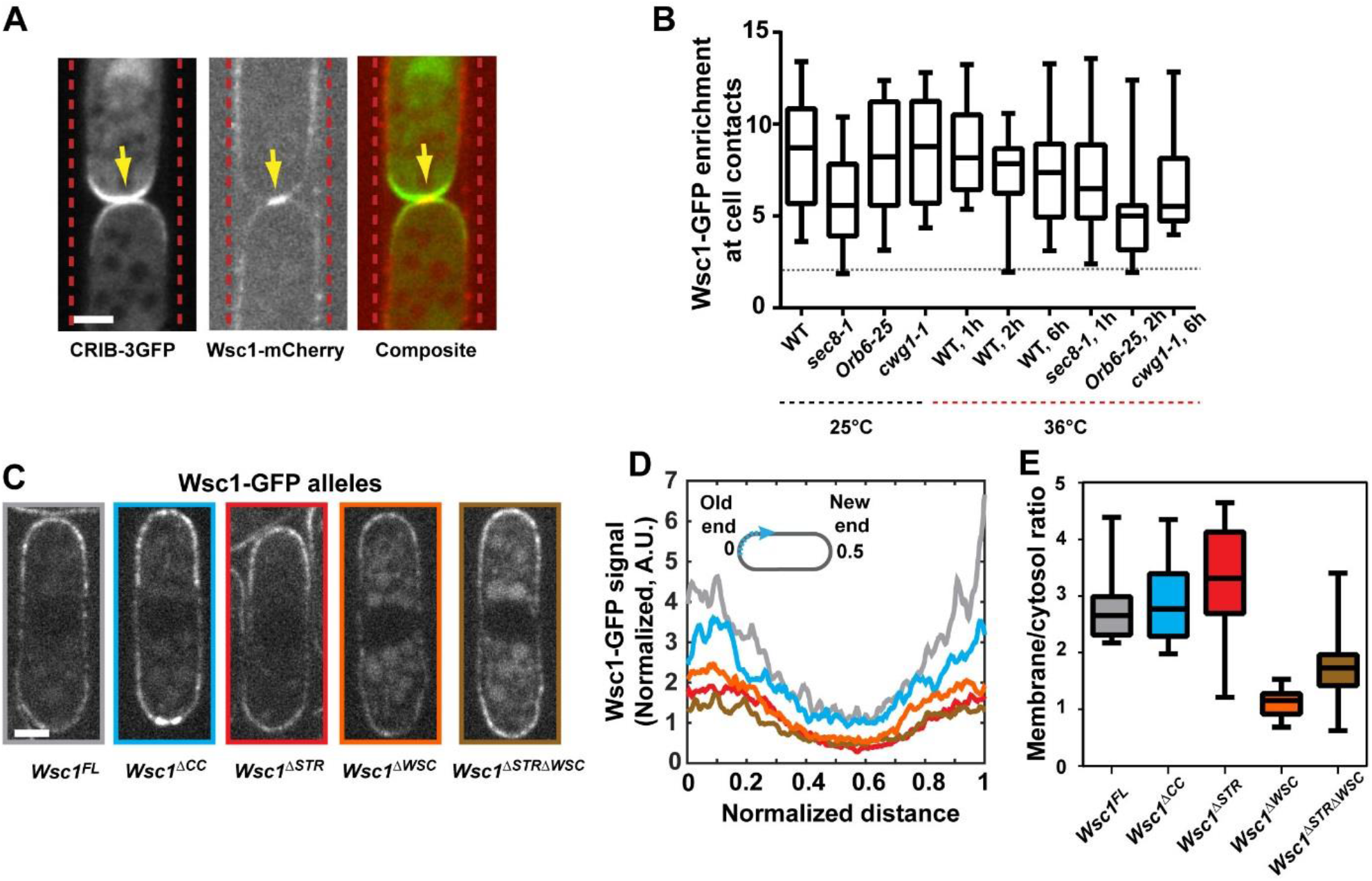
Functional dissection of Wsc1 cluster formation in response to forces onto the Cell Wall. **(A)** Spinning disk confocal mid-slice of WT cells co-expressing CRIB-GFP (a marker for polarity), and Wsc1-mCherry, forming a contact in microchannels **(B)** Quantification of Wsc1-GFP accumulation in the indicated mutants and conditions. The dotted line in the graph correspond to a reference value of enrichment of 2, corresponding to the presence of two apposed membranes. **(C)** Representative mid-slice confocal images of GFP-tagged Wsc1 alleles. **(D)** Normalized fluorescence of GFP signal around cells in different Wsc1 alleles, line colors correspond to boxes colors around images in panel C (n>15 cells for each allele). **(E)** Quantification of membrane/cytoplasm fluorescent signals for all GFP-tagged Wsc1 alleles (n>15 cells for each allele). Scale bars, 2 μm.

**Figure S4(Related to Figure 4).**
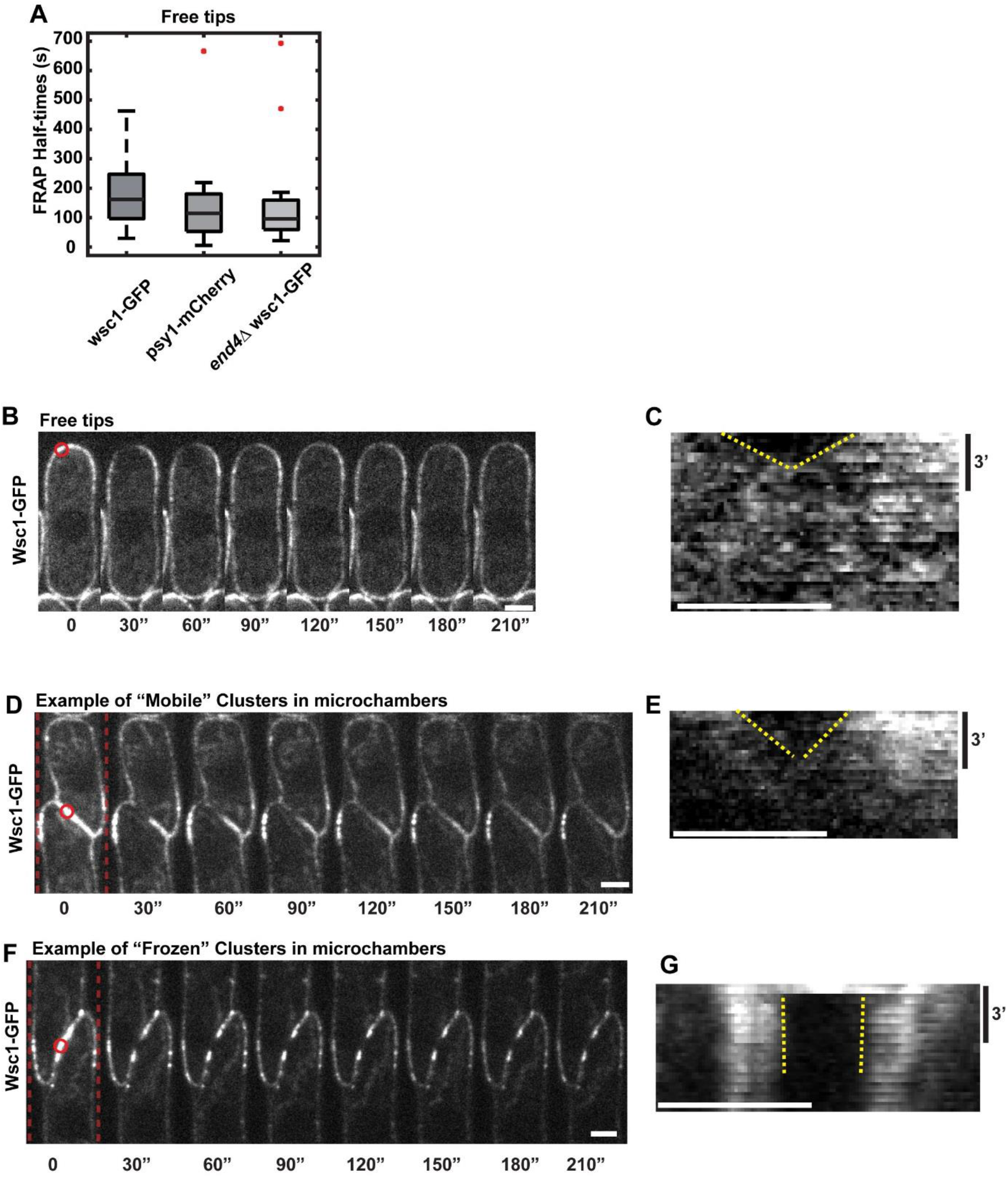
FRAP examples and quantifications. **(A)** FRAP half-times for the indicated markers and mutants, at free growing tips. Red dots are outliers in the distribution (n=27, 12 and 14 cells respectively, from at least 2 independent experiments) **(B)** Representative spinning disk confocal mid-slice time-lapse of a WT cell expressing Wsc1-GFP, where a small region delineated by the red dotted circle was bleached. **(C)** Kymograph corresponding to the experiment presented in panel B computed from a line scan around the cell tip of 3px in width. The yellow dotted lines mark the recovery of the signal in the bleached area. **(D)** Example of diffusing clusters: Spinning disk confocal mid-slice time-lapse of a contact site between WT cells expressing Wsc1-GFP forming a large cluster, where a small region delineated by the red dotted circle was bleached. **(E)** Kymograph corresponding to the experiment presented in panel D computed from a line scan around the cell tip of 3px in width. The yellow dotted lines mark the recovery of the signal in the bleached area. **(D)** Example of frozen clusters: Spinning disk confocal mid-slice time-lapse of a contact site between WT cells expressing Wsc1-GFP forming a large cluster, where a small region delineated by the red dotted circle was bleached. **(E)** Kymograph corresponding to the experiment presented in panel D computed from a line scan around the cell tip of 3px in width. The yellow dotted lines mark the recovery of the signal in the bleached area. Scale bars, 2 μm.

**Figure S5(Related to Figure 5g).**
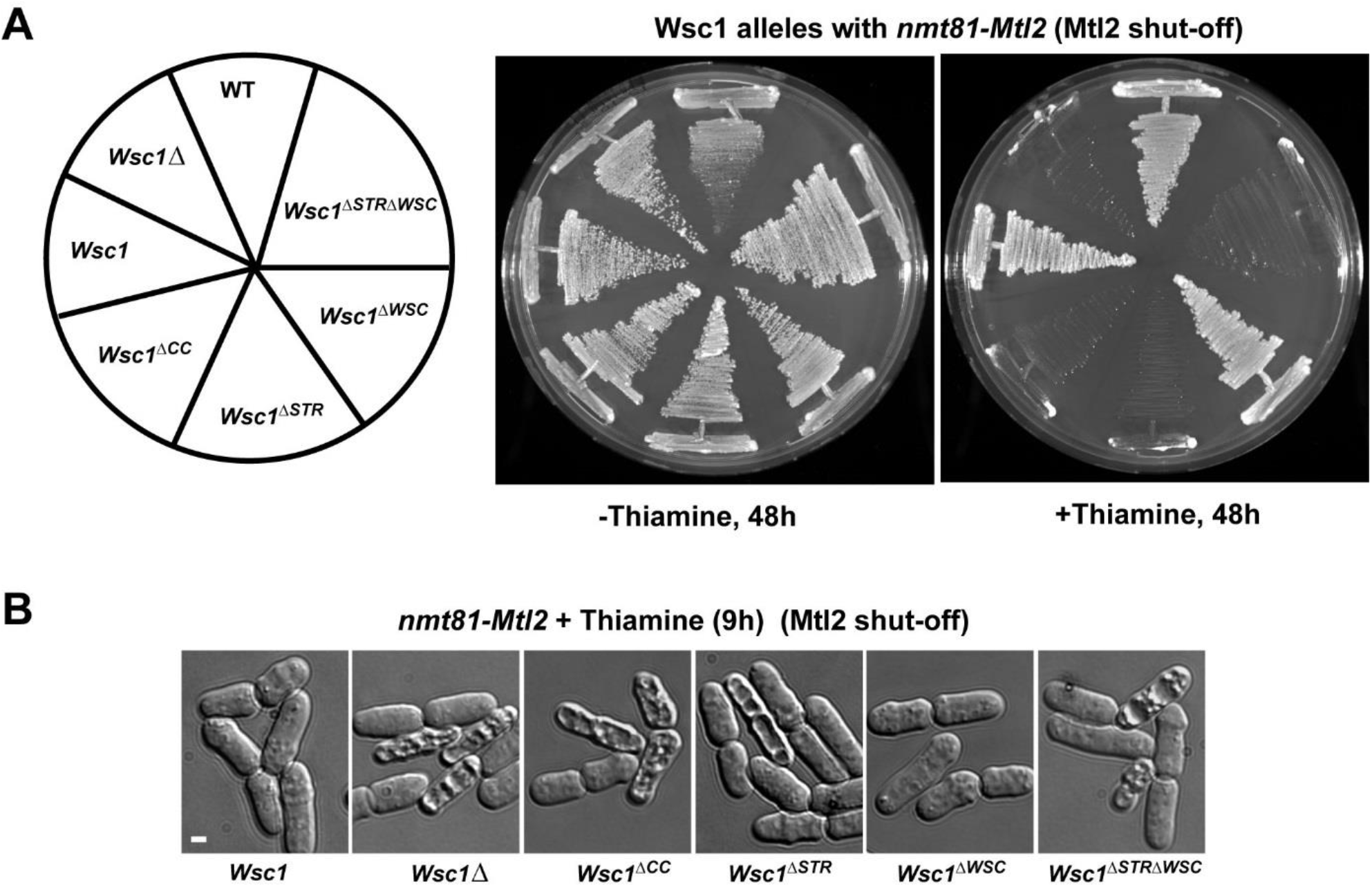
Survival of Wsc1 alleles in the absence of Mtl2. **(A)** Growth for 48h at 25°C of different Wsc1 untagged alleles in a nmt81-Mtl2 background in the absence (Mtl2 expressed); or in the presence of thiamine (Mtl2 shut-off). **(B)** DIC images of Wsc1 untagged alleles in an *nmt81-Mtl2* background grown in liquid cultures in the presence of thiamine for 9h, and placed on agar pads for imaging. Scale bars, 2µm.

**Table S1.** Strains used in this study

|  |  |  |
| --- | --- | --- |
| SM215 | h<sup>-</sup> *leu1-32 ura4D-18* | Sanchez lab stocks |
| GRG45 | h<sup>-</sup> *wsc1-GFP:ura4<sup>+</sup> leu1-32 ade6M210 ura4D-18 his3DI* | Sanchez lab stocks |
| GRG8 | h<sup>-</sup> *wsc1::ura<sup>+</sup> leu1-32 ade6M210 ura4D-18 his3DI* | Sanchez lab stocks |
| SC136 | h<sup>-</sup> *wsc1::his3<sup>+</sup> leu1-32 ade6M210 ura4D-18 his3DI* | Sanchez lab stocks |
| GRG15 | h<sup>-</sup> *wsc1::kan<sup>+</sup> leu1-32 ade6M210 ura4D-18 his3DI* | Sanchez lab stocks |
| YS1220 | h<sup>-</sup> *mtl2::his3 ura4d18 leu1-32 ade6M210 his3DI* | Sanchez lab stocks |
| SC33 | h<sup>-</sup> *kan-81xnm1-mtl2<sup>+</sup> leu1-32 ura4D18 his3D ade6M210* | Sanchez lab stocks |
| SC92 | h<sup>-</sup> *mtl2::kan ura4d18 leu1-32 ade6M210 his3DI* | Sanchez lab stocks |
| SC80 | h<sup>+</sup> *kan-P81xnm1-mtl2<sup>+</sup> wsc1::ura<sup>+</sup> leu1-32 ura4D18 his3D ade6M210* | Sanchez lab stocks |
| SC255 | h<sup>-</sup> *wsc1-GFP:ura4<sup>+</sup> mtl2::kan his3DI leu1-32 ade6M210* | Sanchez lab stocks |
| DB146 | *h90 leu1-32 ade6M210 ura4D-18* | Minc lab stocks |
| RN19 | *h+ wsc1-GFP:ura4<sup>+</sup> leu1-32 ade6M210 ura4D-18* | Minc lab stocks |
| RN73 | *h- wsc1-GFP:ura4<sup>+</sup> leu1-32 ade6M210 ura4D-18* | Minc lab stocks |
| RN21 | *h- mtl2-GFP:ura4<sup>+</sup> leu1-32 ade6M210 ura4D-18* | Minc lab stocks |
| RN42 | *h90 wsc1-GFP:ura4<sup>+</sup> leu1-32 ura4D-18 ade-* | This study |
| RN48 | *h+ wsc1-mCherry:KanMX leu1-32 ade6M210 ura4D-18* | This study |
| RN51 | *h90 wsc1:: KanMX leu1-32 ade6M210 ura4D-18* | This study |
| RN46 | *h90 wsc1-GFP:ura4<sup>+</sup> leu:mCherry-Psy1 ura4D-18 ade-* | This study |
| RN57 | *h90 wsc1-mCherry:KanMX mtl2-GFP:ura4<sup>+</sup>* | This study |
| RN126 | *h90 wsc1-GFP:ura4<sup>+</sup> gap1::hphMX* | This study |
| RN127 | *h90 wsc1-GFP:ura4<sup>+</sup> fus1:: KanMX* | This study |
| NM448 | *h+ leu:mCherry-psy1 ura4-D18 ade-* | Lab stocks |
| NM447 | *h90 leu:mCherry-psy1 ura4-D18 ade-* | Lab stocks |
| RN170 | *h90 wsc1-GFP:ura4<sup>+</sup> ade6+:ptdh1:3mTagBFP2-Ritc:terminatorScADH1:bsdMX* | This study |
| RN54 | *h90 wsc1-mCherry:KanMX CRIB-3GFP:ura leu1-32 ura4-D18* | This study |
| RN91 | *h90 wsc1-GFP:ura4<sup>+</sup> bgs4::ura4 RFP-bgs4-Leu ade-* | This study |
| RN82 | *h90 wsc1-GFP:ura4<sup>+</sup> end4::KanMX leu1-32 ura4-D18 ade-* | This study |
| RN90 | *h90 wsc1-GFP:ura4<sup>+</sup> wsp1::leu ade-* | This study |
| RN88 | *h90 wsc1-GFP:ura4<sup>+</sup> exo70::KanMX leu1-32 ura4-D18 ade-* | This study |
| RN152 | *h90 wsc1-GFP:ura4<sup>+</sup> sec8-1 ade6 Leu1-32 ura4-D18* | This study |
| RN84 | *h90 wsc1-GFP:ura4<sup>+</sup> for3::KanMX* | This study |
| RN138 | *h90 wsc1-GFP:ura4<sup>+</sup> myo52::KanMX* | This study |
| RN107 | *h90 wsc1-GFP:ura4<sup>+</sup> orb6-25 ade6-M210 leu1-32* | This study |
| RN153 | *h90 wsc1-GFP:ura4<sup>+</sup> cwg 1-1* | This study |
| RN77 | *h90 wsc1-GFP:ura4<sup>+</sup> pck1::KanMX leu1-32 ura4-D18ade-* | This study |
| RN76 | *h90 wsc1-GFP:ura4<sup>+</sup> pmk1::KanMX leu1-32 ura4-D18 ade-* | This study |
| RN143 | *h90 wsc1-GFP:ura4<sup>+</sup> styl::kanMX leu1-32 ura4-D18 ade-* | This study |
| RN109 | *h90 wsc1-GFP:ura4<sup>+</sup> rgf1::KanMX leu1-32 ura4-D18 ade-* | This study |
| RN86 | *h90 wsc1-GFP:ura4<sup>+</sup> rgf2::KanMX leu1-32 ura4-D18 ade-* | This study |
| RN129 | *h90 wsc1-GFP:ura4<sup>+</sup> mtl2::KanMX leu1-32 ura4-D18 ade-* | This study |
| RC90 | h<sup>-</sup> *wsc1<sup>ΔWSC</sup>-GFP:ura4<sup>+</sup> leu1-32 ade6M210 ura4D-18 his3DI* | This study |
| RC96 | h<sup>-</sup> *wsc1<sup>ΔSTR</sup>-GFP:ura4<sup>+</sup> leu1-32 ade6M210 ura4D-18 his3DI* | This study |
| RC98 | h<sup>-</sup> *wsc1<sup>ΔSTRΔWSC</sup>-GFP:ura4<sup>+</sup> leu1-32 ade6M210 ura4D-18 his3DI* | This study |
| RC91 | h<sup>-</sup> *wsc1<sup>ΔCC</sup>-GFP:ura4<sup>+</sup> leu1-32 ade6M210 ura4D-18 his3DI* | This study |
| RN116 | *h90 wsc1<sup>ΔCC</sup>-GFP:ura4<sup>+</sup> leu1-32 ade6M210 ura4D-18* | This study |
| RN122 | *h90 wsc1<sup>ΔSTR</sup>-GFP:ura4<sup>+</sup> leu1-32 ade6M210 ura4D-18* | This study |
| RN112 | *h90 wsc1<sup>ΔWSC</sup>-GFP:ura4<sup>+</sup> leu1-32 ade6M210 ura4D-18* | This study |
| RN125 | *h90 wsc1<sup>ΔSTRΔWSC</sup>-GFP:ura4<sup>+</sup> leu1-32 ade6M210 ura4D-18* | This study |
| RN62 | *h90 wsc1-mCherry:KanMX leu1:rgf1-GFP* | This study |
| RN63 | *h90 wsc1-mCherry:NatMX leu1::KanMX-P3nmt1-pkc1(HR1-C2)-mECitrine* | This study |
| RN145 | *h90 wsc1-mCherry:NatMX pck1::pck1-GFP-HA-KanMX* | This study |
| RN146 | *h90 wsc1-mCherry:NatMX pck2::pck2-GFP-HA-KanMX* | This study |
| RN176 | *h90 wsc1<sup>FL</sup>:ura4<sup>+</sup> leu1-32 ade6M210 ura4D-18* | This study |
| RN178 | *h90 wsc1<sup>ΔCC</sup>:ura4<sup>+</sup> leu1-32 ade6M210 ura4D-18* | This study |
| RN182 | *h90 wsc1<sup>ΔSTR</sup>:ura4<sup>+</sup> leu1-32 ade6M210 ura4D-18* | This study |
| RN180 | *h90 wsc1<sup>ΔWSC</sup>:ura4<sup>+</sup> leu1-32 ade6M210 ura4D-18* | This study |
| RN184 | *h90 wsc1<sup>ΔSTRΔWSC</sup>:ura4<sup>+</sup> leu1-32 ade6M210 ura4D-18* | This study |
| RN221 | *h90 wsc1<sup>ΔSTR MTL2-STR</sup>-GFP:ura4<sup>+</sup> leu1-32 ura4D-18* | This study |
| YS5505 | h<sup>+</sup> *wsc1<sup>ΔWSC</sup>:ura4<sup>+</sup> kan-P81xnm1-mtl2<sup>+</sup> leu1-32 ura4D18 ade6M210* | This study |
| YS5485 | h<sup>+</sup> *wsc1<sup>ΔWSC ΔSTR</sup>:ura4<sup>+</sup> kan-P81xnm1-mtl2<sup>+</sup> leu1-32 ura4D18 ade6M210* | This study |
| YS5504 | h<sup>+</sup> *wsc1<sup>ΔCC</sup>:ura4<sup>+</sup> kan-P81xnm1-mtl2<sup>+</sup> leu1-32 ura4D18 ade6M210* | This study |
| YS5509 | h<sup>-</sup> *wsc1<sup>ΔSTR MTL2-STR</sup>-GFP::ura4<sup>+</sup> leu1-32 ade6M210 ura4D-18* | This study |

## SUPPLEMENTAL MOVIE LEGENDS

**Movie S1 (related to Figure 1). Wsc1 clusters at sites of septation**. Example of dividing Wsc1-GFP cells forming a firm contact upon septation, driving Wsc1 sensor accumulation. Time is in hh:min.

**Movie S2 (Related to Figure 1). Wsc1 clusters dissolve when cells slide past each other**. Fields of dividing and growing cells that form clusters at new ends. These Wsc1 clusters disappear as cells slide past each other’s. Time is in hh:min.

**Movie S3 (Related to Figure 1). Wsc1 clusters at old end old end contacts**. Example of growing Wsc1-GFP cells that contact at old ends forming a firm contact at old ends, yielding the formation of a bright Wsc1 cluster. Time is in hh:min.

**Movie S4 (Related to Figure 1). Wsc1 clusters at end contacts in microchannels**. Example of growing Wsc1-GFP cells confined in a linear microchannel that contact at cell tips, yielding the formation of bright and stable Wsc1 clusters at end contacts. Time is in hh:min.

**Movie S5 (Related to Figure 1). Wsc1 clusters at mating tip contacts**. Example of mating Wsc1-GFP cells that for mating tips that grow on each other and form a firm contact at mating tips, yielding the formation of a bright Wsc1 cluster. Time is in hh:min.

